# Fitness Factor Genes Conserved within the Multi-species Core Genome of Gram-negative Enterobacterales Species Contribute to Bacteremia Pathogenesis

**DOI:** 10.1101/2024.03.18.585282

**Authors:** Harry L. T. Mobley, Mark T. Anderson, Bridget S. Moricz, Geoffrey B. Severin, Caitlyn L. Holmes, Elizabeth N. Ottosen, Tad Eichler, Surbhi Gupta, Santosh Paudel, Ritam Sinha, Sophia Mason, Stephanie D. Himpsl, Aric N. Brown, Margaret Gaca, Christina M. Kiser, Thomas H. Clarke, Derrick E. Fouts, Victor J. DiRita, Michael A. Bachman

## Abstract

There is a critical gap in knowledge about how Gram-negative bacterial pathogens, using survival strategies developed for other niches, cause lethal bacteremia. Facultative anaerobic species of the Enterobacterales order are the most common cause of Gram-negative bacteremia, including *Escherichia coli*, *Klebsiella pneumoniae*, *Serratia marcescens, Citrobacter freundii,* and *Enterobacter hormaechei*. Bacteremia often leads to sepsis, a life-threatening organ dysfunction resulting from an unregulated immune response to infection. Despite a lack of specialization for this host environment, Gram-negative pathogens cause nearly half of bacteremia cases annually. Based on our existing Tn-Seq fitness factor data from a murine model of bacteremia combined with comparative genomics of the five Enterobacterales species above, we prioritized 18 conserved fitness genes or operons for further characterization. Each mutant in each species was used to cochallenge C57BL/6 mice via tail vein injection along with the respective wild-type strain to determine competitive indices for each fitness gene or operon. Among the five species, we found three fitness factor genes, that when mutated, attenuated the mutant for all species in the spleen and liver (*tatC, ruvA, gmhB*). Nine additional fitness factor genes or operons were validated as outcompeted by wild-type in three or four bacterial species in the spleen (*xerC*, *wzxE*, *arcA*, *prc*, *apaGH*, *atpG*, *lpdA*, *ubiH*, *aroC*). Overall, 17 of 18 fitness factor mutants were attenuated in at least one species in the spleen or liver. Together, these findings allow for the development of a model of bacteremia pathogenesis that may include future targets of therapy against bloodstream infections.

**>Author Summary:** Frequent cases of bacteremia plague our ICUs, bone marrow transplant units, and inpatient facilities. Nearly half of these infections are caused by Gram-negative bacteria. The Enterobacterales order including *E. coli*, *K. pneumoniae, S. marcescens, C. freundii*, and *E. hormaechei* are leading causes of bacteremia. An alarming proportion of these are due to antibiotic-resistant isolates, which are four times more likely to kill than antibiotic-susceptible isolates. Clearly, we need new therapeutic targets to treat cases of bacteremia and sepsis. Previously, it has been unclear what genes contribute to their ability to survive in this hostile host environment. We have previously undertaken unbiased genetic screens to identify 18 genes shared by all five bacterial genera that are required for survival in blood and blood-filtering organs. These include genes that encode proteins that maintain proton motive force, resist antimicrobial peptides and complement, mediate genome maintenance, transport key metabolites and proteins, avoid oxidative stress, acquire iron, and regulate key pathways. Mutants, constructed in these shared genes in the five species, were validated for a high proportion of genes as critical for infection in the mouse model of bacteremia.

## Introduction

Sepsis is life-threatening organ dysfunction that results from an unregulated immune response to infection. It is the leading cause of death in hospitalized patients across the United States (1) with a mortality rate of 25-50% leading to 220,000 deaths per year. Bacteremia is a leading cause of sepsis and Gram-negative pathogens cause nearly half of bacteremia cases annually (2). Species within the Enterobacterales order are the most common cause of Gram-negative bacteremia, including the species *Escherichia coli*, *Klebsiella pneumoniae*, *Serratia marcescens, Citrobacter freundii* (3) and *Enterobacter hormaechei* (4). Early treatment with antibiotics is critical to reduce mortality, but antibiotic resistance may thwart this empiric therapy. There is a critical need to develop new therapies and salvage existing ones, so that we can counter antibiotic resistance and reduce sepsis mortality.

Bacteremia has three phases of pathogenesis: initial primary site infection, dissemination to the bloodstream, and growth and survival in blood and blood-filtering organs (5). In Gram-negative bacteremia, the primary site serves as a reservoir of the pathogen that can intermittently re-seed the bloodstream and prolong the infection. We have recently determined that Enterobacterales replicate rapidly in the liver and spleen during bacteremia (6), but are slowly cleared in most cases, indicating that the immune system can overcome rapid bacterial growth. Whereas current antibiotics are based on the ability to kill or inhibit bacterial growth *in vitro*, there is an opportunity to identify drug targets that are specifically required during infection.

To that end, we previously constructed random transposon libraries or ordered transposon libraries in representative clinical isolates of five Gram-negative Enterobacterales species: *E. coli* CFT073 (7), *K. pneumoniae* KPPR1 (8, 9), *S. marcescens* UMH9 (10), *C. freundii* UMH14 (11), and *E. hormaechei* UM_CRE_14 (Ottosen, in preparation). We conducted global Tn-Seq screens in the murine model of bacteremia by tail vein inoculation and identified putative fitness genes predicted to be required by each species to survive in the bloodstream and colonize the spleen. The characteristics of the Tn libraries and the outcomes of the Tn-Seq screens are summarized in **Table 1**.

**Table 1.** Species, strains, transposon pool complexity, and genome sequences used in Tn-Seq studies in the murine bacteremia model.

| Species and strain | No. mutants in transposon pool | No. fitness genes predicted by Tn-Seq | Fitness genes validated<sup>a</sup> | Genome size (Mb) (species range) | <u>Pan genome analysis</u> |  | Reference |
| --- | --- | --- | --- | --- | --- | --- | --- |
|  |  |  |  |  | No. genomes sequenced by us | No. genomes in global database |  |
| *E. coli* CFT073 | 360,000 | 242 | 9/11 | 5.2 (4.9-5.4) | 27 | 9904 | (7) |
| *K. pneumoniae* KPPR1 | 25,000 | 58 | 8/8 | 5.4 (4.2-6.1) | 117 | 3517 | (12)<br>(13)<br>(8) |
| *C. freundii* UMH14 | 44,000 | 223 | 5/7 | 4.9 (4.9-5.8) | 8 | 102 | (11) |
| *S. marcescens* UMH9 | 32,000 | 169 | 6/7 | 5.0 (4.9-5.4) | 11 | 358 | (10) |
| *E. hormaechei* UM_CRE_14 | 15,000 | 214 | 5/7 | 5.0 (5.1-5.6) | 90 | 593 | (Ottosen *et al.*, in preparation) |

<sup>a</sup>number of independently constructed fitness factor mutants that were validated in murine model of bacteremia per number of fitness factors tested in the original published or submitted studies for each species.

To further enable these studies, we also conducted extensive genomic comparisons and identified the multi-species core genome of Enterobacterales species commonly causing bacteremia in humans (Fouts, Derrick E., Thomas H. Clarke, Aric N. Brown, Elizabeth N. Ottosen, Caitlyn Holmes, Bridget S. Moricz, Sophia Mason, Ritam Sinha, Geoffrey B. Severin, Mark T. Anderson, Victor DiRita, Michael A. Bachman and Harry L. T. Mobley Integration of Pan-genome with Operon Organization, Virulence factors, Antimicrobial resistance and Tn-Seq of Gram-negative *Enterobacterales* Commonly Causing Bacteremia (submitted). By integrating our pan-genome and genome-wide fitness data, we aligned Tn-Seq screen hits identified in a murine model of bacteremia to predicted fitness genes shared among Enterobacterales species.

Although phenotypically similar in terms of antimicrobial resistance and biochemical identification tests, these five Gram-negative species nevertheless represent a heterogeneous group of organisms that differ in virulence mechanisms, primary sites of infection, and metabolic pathways. There is also wide variation in our knowledge regarding infections of the bloodstream. For example, *E. coli* has been extensively studied in the context of extraintestinal infection (∼6000 PubMed references on “*E. coli* and bacteremia” as of 2023) and ∼2000 for *K. pneumoniae.* In contrast, our understanding of *S. marcescens, C. freundii,* and *E. hormaechei* has lagged behind despite the recognition of their epidemiological importance (just over 300 references combined). Moreover, only a small percentage of these reports deal with virulence mechanisms, but rather characterize clinical infections and antibiotic resistance. While previous studies have evaluated Gram-negative bacteremia fitness genes required in the blood, there has been no systematic analysis of shared genes critical across species in this hostile environment. Here, we address the relative lack of rigorous studies of the pathogenesis and potential for novel treatments for Enterobacterales bacteremia.

The goal of this study was to identify and characterize conserved bacterial fitness genes and operons that play critical roles in the development and outcome of bacteremia that may also serve as potential therapeutic targets. Data from independent Tn-Seq studies of five Enterobacterales species in a murine model of bacteremia were used to prioritize conserved fitness factor genetic components. Mutants in 18 shared genetic components were constructed in all five species and used to cochallenge C57BL/6 mice via tail vein injection along with the respective wild-type strain of each species to determine competitive indices for each fitness gene or operon. Twelve of 18 prioritized components were confirmed as fitness factors in three or more species. Relevant phenotypes of the mutants were assessed to validate the mutant constructs and to identify potential *in vitro* correlates of virulence. We propose that these genes and their respective proteins may represent future targets of therapy against bloodstream infections.

## Results

### Selection and mutation of fitness genes found in the multi-species core genome shared by at least four of the five species

In preparation for this study, we defined a “multi-species core genome” composed of 2850 genes shared by at least four of the five species at a minimum of 70% predicted protein identity (Fouts *et al*., submitted). Using the Tn-Seq data from transposon libraries of the Gram-negative species used to challenge our murine model via tail injection, we found 500 total bacteremia fitness genes present in at least one species-level core-genome of which 373 were represented in the multi-species core genome (*i.e.*, present in all five species-level core-genomes). From the 500 total bacteremia fitness genes mentioned above, we created a scoring rubric to prioritize conserved fitness genes of interest for further study based on the following criteria (Fouts *et al*., submitted); 1) the magnitude of the fitness defect associated with a gene in any one species, 2) a gene that was a fitness factor in multiple species, 3) whether multiple fitness genes reside in the same operon, and 4) that mutation of a fitness gene was found to confer increased antibiotic susceptibility in *E. coli* BW25113 (14, 15). 102 of the 500 fitness factor genes met this last criterion, including *atpG, ubiG, hscB, gmhB, sapA, pstC, tatC, ruvA, xerC* and *lpdA*. Overall, 18 single genes and/or gene clusters within operons, ranked according to the scoring rubric, were prioritized for study based on their initial scoring (**Table 2**). These conserved and shared bacteremia fitness genes, predicted by Tn-Seq screens as required for virulence of Gram-negative Enterobacterales bacilli, were assigned to seven common pathways.

**Table 2.** Prioritization criteria for conserved and shared bacteremia fitness genes within common pathways predicted by Tn-Seq screens as required for virulence of Gram-negative Enterobacterales bacilli.

| Bacterial Cell Function | Scoring Rubric Score |  |  |  | Phenotypes of mutants assessed |
| --- | --- | --- | --- | --- | --- |
| 1. Maintenance of Proton-Motive Force across inner membrane | Gene<sup>a</sup> | Score | Operon<sup>b</sup> | Score |  |
| ATP Synthase | *atpG* | 9 | *atpIBEFHAGDC* | 40 | + |
| Electron Transport: Ubiquinone synthesis | *ubiH* | 5 | *ygfB pepP ubiHI* | 14 |  |
| Iron-Sulfur Cluster biosynthesis for cytochrome maturation | *hscB* | 10 | *iscRSUA hscBA fdx iscX* | 36 |  |
| 2. Resistance to Antimicrobial Peptides and Complement |  |  |  |  |  |
| Modification of Lipid A of LPS | *arnD* | 12 | *arnBCADTEF* | 34 | + |
| LPS inner core synthesis | *gmhB* | 7 | *gmhB* | n/a<sup>c</sup> | + |
| Synthesis of Enterobacterial Common Antigen | *wzxE* | 13 | *wecA wzzE wecBC rffGH wecDE wzxE wecF wzyE rffM* | 59 | + |
| Sensitivity to antimicrobial peptides | *sapA* | 8 | *sapABCD* | 19 | + |
| Periplasmic protease inactivates complement | *prc* | 6 | *proQ prc* | 16 | + |
| 3. Transport |  |  |  |  |  |
| Phosphate (ABC) transport | *pstC* | 10 | *pstSCABphoU* | 31 | + |
| Twin arginine transporter | *tatC* | 9 | *tatABCD* | 11 | + |
| **4. Genome maintenance** |  |  |  |  |  |
| Endonuclease that resolves Holliday junctions | *ruvA* | 10 | *ruvAB* | 12 | + |
| Tyrosine recombinase | *xerC* | 10 | *yifL dapF yigA xerC yigB* | 24 | + |
| **5. Shikimate biosynthesis** |  |  |  |  |  |
| Contributes chorismate for quinone, siderophore, aromatic amino acid, and folate biosynthesis | *aroK<sup>e</sup>* | 6 | *aroKB damX dam rpe gph trpS* | 25 | + |
| **6. Global gene regulation** |  |  |  |  |  |
| Aerobic respiration control protein | *arcA* | 5 | *arcA* | n/a<sup>c</sup> | + |
| **7. Osmotic and Oxidative stress** |  |  |  |  |  |
| Regulates Import of proline/betaine by ProP | *proQ<sup>d</sup>* | 10 | *proQ prc* | 16 | + |
| Oxidative stress response | *apaG* | 4 | *lptD surA pdxA rsmA apaGH* | 31 | + |
|  | *pdxA* | 8 | (same as above) | 31 |  |
| **8. Metabolic Pathways** |  |  |  |  |  |
| Pyruvate dehydrogenase complex component | *lpdA* | 10 | *pdhR aceEF lpdA* | 17 |  |

<sup>a</sup>individual gene score from scoring rubric (Fouts *et al.*, submitted)
<sup>b</sup>score from the scoring rubric of the operon containing the highlighted gene(s); **bold** indicates that a gene was identified as a fitness factor in at least one of the five bacterial species in Tn-Seq studies (**Table 1**). Operon gene order is representative for the five species, however, gene order may differ in some species.
<sup>c</sup>n/a, not applicable; gene not found within an operon
<sup>d</sup>because of concerns that a *proQ* mutant would introduce polar effects on the downstream *prc* gene, *proP*, which encode the proline/betaine importer that is regulated by *proQ*, was chosen to construct the mutant used in this study.
<sup>e</sup>the *aroC* allele, chorismate synthase, encodes the terminal enzyme in the shikimate biosynthesis pathway and was selected for mutagenesis in this study.

All isolates representing the five species (**Table 1**) have been demonstrated as amenable to *lambda* red recombineering (see “Construction of mutants in prioritized fitness genes” in Materials and Methods). Therefore, using a common strategy outlined in **Fig 1**, we constructed 89 total mutants composed of either single or multi-gene mutations (*i.e.*, operons) covering 203 genes in total across all five species. In *E. coli K. pneumoniae, S. marcescens,* and *C. freundii*, we constructed 18 mutants that covered 42 total genes per species. In *E. hormaechei* UM_CRE_14, a strain that lacks *arn* genes, we constructed 17 mutants that covered 35 genes. All mutant constructs were verified by PCR and/or sequencing.

**Figure 1.**
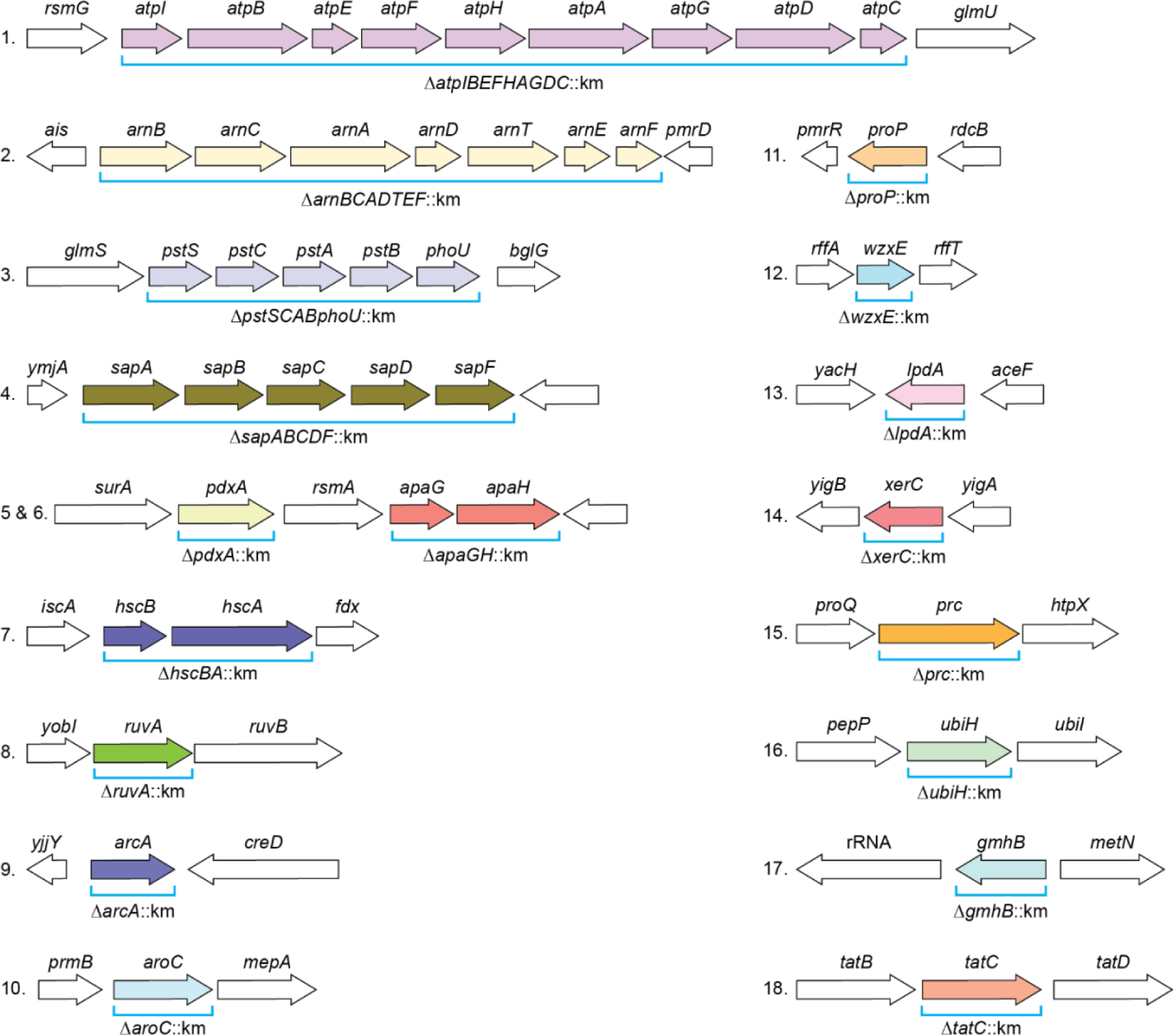
Prioritized multi-species bacteremia fitness genes. Colored arrows represent fitness genes that were targeted for mutagenesis via recombineering. The genetic organization of *E. coli* CFT073 is shown for reference, but corresponding mutations were generated in each of the five species of interest. ORF lengths are not to scale.

### Assessment of virulence of mutants versus wildtype strains in the murine tail vein injection model

Each of the 89 mutants were competed head-to-head against their respective wild-type strains in co-infections in the murine bacteremia model. At 24 hours post-inoculation, mice were euthanized and CFU/g tissue in homogenates of spleens and livers were enumerated and used to calculate competitive indices. Results are shown for all five species in **Table 3** for spleens and **Table 4** for livers. We additionally report competitive indices of *S. marcescens* UMH9 mutants in kidneys (**Table 5**) owing to the wild-type’s robust capacity to colonize this organ in this murine model (6). Individual mutant competitive indices found to be statistically below the hypothetical mean of 0 (*i.e*., neutral fitness) are considered attenuated (green shading). Competitive indices were derived from pools of 5-14 mice.

**Table 3.** Competitive indices of bacteremia Tn-Seq fitness factor gene mutants in the <u>Spleen</u> following cochallenge via tail vein injection with respective wild-type parent strain in the murine model of bacteremia.

| Prioritized fitness factor gene | *E. coli* CFT073 | *K. pneumoniae* KPPR1 | *C. freundii* UMH14 | *S. marcescens* UMH9 | *E. hormaechei* UM_CRE_14 |
| --- | --- | --- | --- | --- | --- |
|  | Log<sub>10</sub> Competitive Index<sup>a</sup> (*p* value<sup>b</sup>)<sup>c</sup> |  |  |  |  |
| *tatC* | -0.440<sup>*d</sup> | -1.725<sup>**</sup> | -1.772<sup>***g</sup> | -2.187<sup>***</sup> | -0.843<sup>***</sup> |
| *ruvA* | -1.191<sup>*</sup> | -1.331<sup>***</sup> | -1.019<sup>***</sup> | -0.675<sup>***</sup> | -1.533<sup>*</sup> |
| *gmhB* | -0.848<sup>***e</sup> | -0.800<sup>***e</sup> | -0.293<sup>***e</sup> | -0.954<sup>***</sup> | -1.326<sup>***</sup> |
| *xerC* | -0.438<sup>**</sup> | -0.747<sup>**</sup> | -0.533<sup>***</sup> | -0.556<sup>*</sup> | -1.512<sup>*</sup> |
| *wzxE* | -0.903<sup>*</sup> | -1.105<sup>***</sup> | 0.063 | -1.077<sup>***</sup> | -0.750<sup>***</sup> |
| *arcA* | -0.154<sup>f</sup> | -0.697<sup>**f</sup> | -0.407<sup>*f</sup> | -0.279<sup>*f</sup> | -1.046<sup>***</sup> |
| *prc* | -0.482<sup>**</sup> | 0.730<sup>*</sup> | -0.222<sup>***</sup> | -0.580<sup>***</sup> | -0.395<sup>**</sup> |
| *apaGH* | 0.058 | -0.519<sup>**</sup> | -0.349<sup>**</sup> | -0.261<sup>*</sup> | -0.518<sup>*</sup> |
| *atpIBEFHAGDC* | -0.6306<sup>**</sup> | -1.531<sup>**</sup> | 0.140 | -0.566<sup>***</sup> | -0.506 |
| *lpdA* | -0.824 | -0.674 | -0.251<sup>*</sup> | -0.557<sup>*</sup> | -1.093<sup>*</sup> |
| *ubiH* | -0.038 | -0.494<sup>***</sup> | -0.093 | -0.649<sup>*</sup> | -0.883<sup>***</sup> |
| *aroC* | -0.018 | -0.092 | -0.564<sup>***</sup> | -0.351<sup>***</sup> | -0.519<sup>**</sup> |
| *pdxA* | -0.261 | -0.356**i | -0.118 | -0.084 | -0.392* |
| *hscBA* | -0.299*** | -0.377** | 0.570 | -0.032 | 0.288 |
| *pstSCABphoU* | -0.387* | -0.031 | -0.0118 | -0.104 | -0.568* |
| *sapABCDF* | 0.015 | -0.924* | 0.032 | 0.391 | -0.222 |
| *arnBCADTEF* | -0.275 | 0.060 | -0.235 | -0.0445 | -<sup>h</sup> |
| *proP* | -0.057 | 0.170* | 0.174* | 0.0048 | 0.060 |

<sup>a</sup>Competitive index (CI) was calculated as $(CFU_{mutant}/CFU_{wild-type})^{output}/(CFU_{mutant}/CFU_{wild-type})^{input}$ . Inocula in total CFU were *E. coli* CFT073 ( $1 \times 10^7$ ), *K. pneumoniae* KPPR1 ( $1 \times 10^5$ ), *S. marcescens* UMH9 ( $5 \times 10^6$ ), *C. freundii* UMH14 ( $1 \times 10^8$ ), and *E. hormaechei* UM\_CRE\_14 ( $1 \times 10^8$ ).
<sup>b</sup>CI data subjected to *t*-test for determination of statistical significance.
\* $p \leq .05$ , \*\* $p \leq .01$ , \*\*\* $p \leq .001$
<sup>c</sup>number of mice co-challenged by tail vein injection with wild-type and mutant were 5-14 mice. CFU determined at 24 hours
<sup>d</sup>Green shading, wild-type significantly outcompeted mutant; No shading, wild-type did not significantly outcompete mutant. Brown shading, mutant significantly outcompeted wild-type strain. Genes are listed in order of number of species in which the gene was validated as a fitness gene and then by the magnitude of fitness defect in validated genes.
<sup>e</sup>*E. coli gmhB* competitive index derived from *gmhB*:Tn5 library (13). *K. pneumoniae* and *C. freundii gmhB* competitive indices have been reported in our published work (11).
<sup>f</sup>*arcA* competitive indices have been reported in our published work (16).
<sup>g</sup>*tatC* competitive indices have been reported in our published work (11).
<sup>h</sup>*arn* genes not present in *E. hormaechei*
<sup>i</sup>*pdxA* competitive index for *K. pneumoniae* has been reported in our published work (8).

**Table 4.**
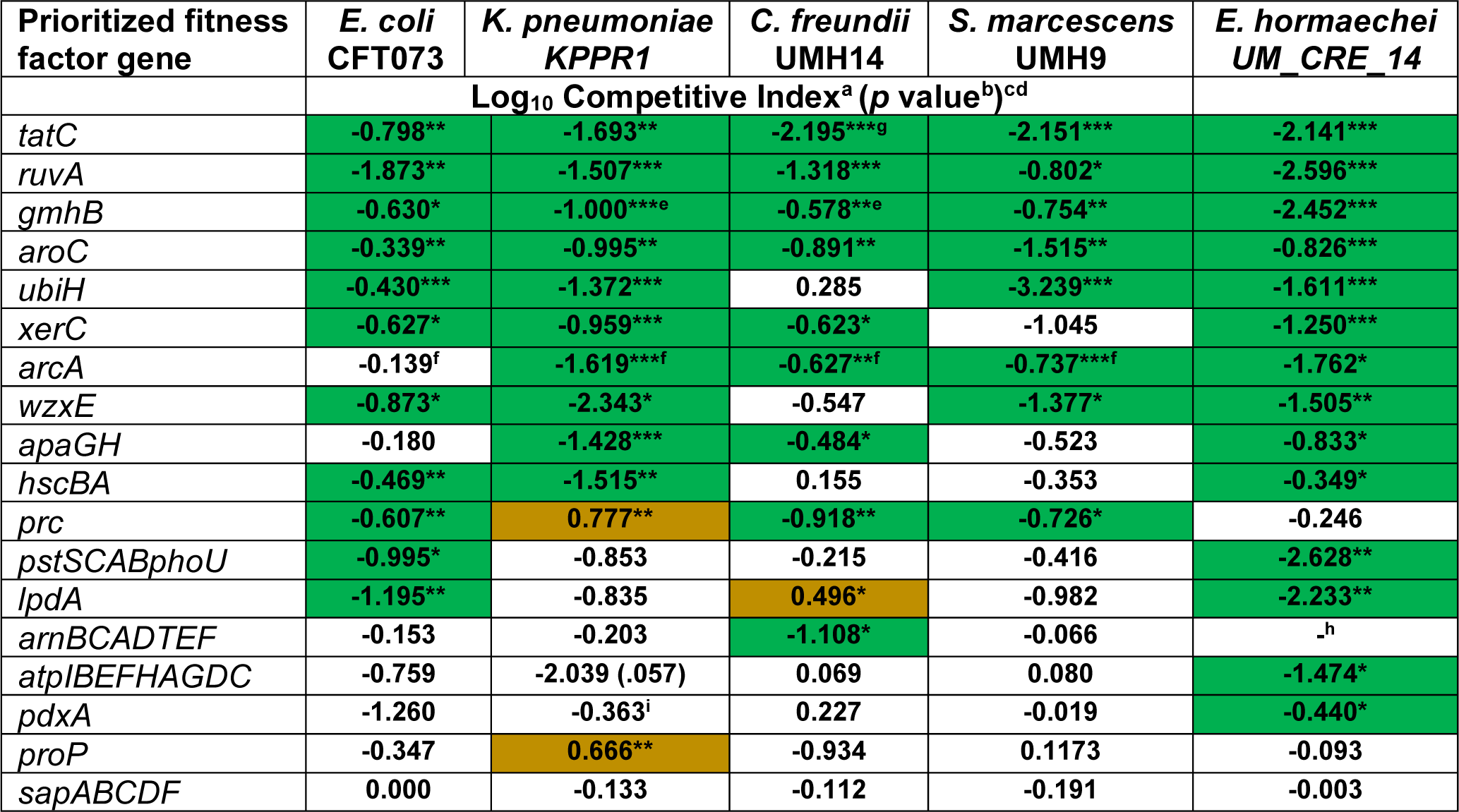
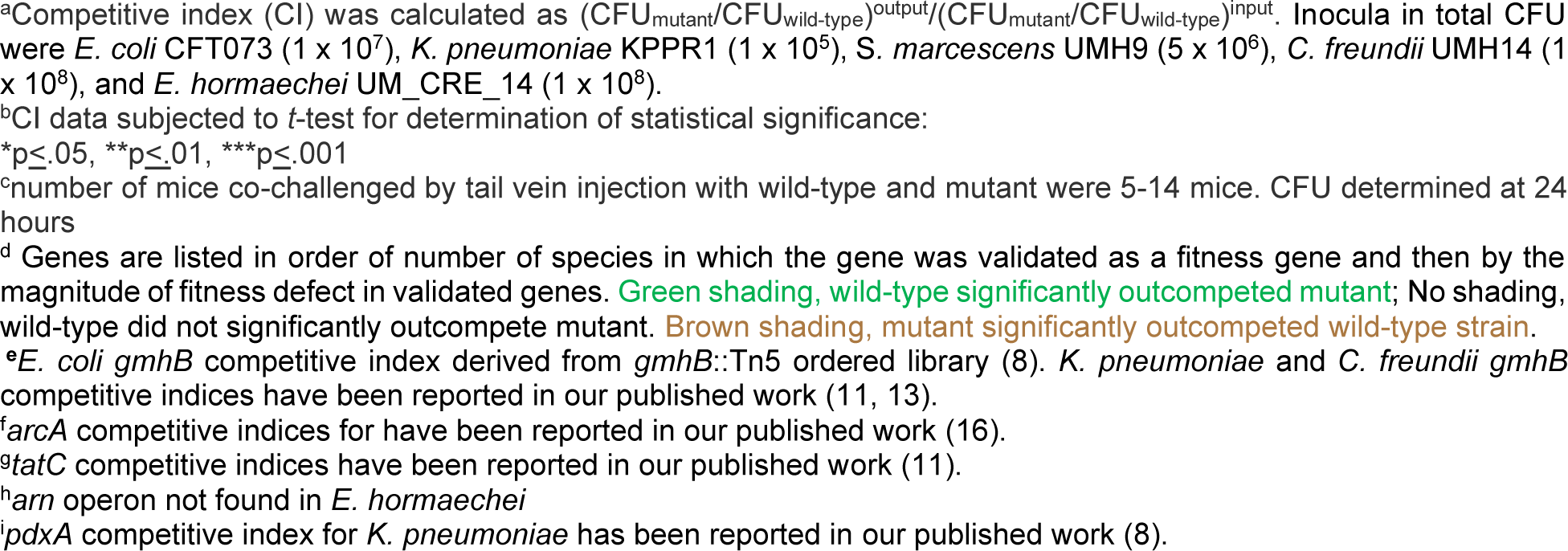
Competitive indices of bacteremia Tn-Seq fitness factor gene mutants in the <u>Liver</u> following cochallenge via tail vein injection with respective wild-type parent strain in the murine model of bacteremia.

**Table 5.**
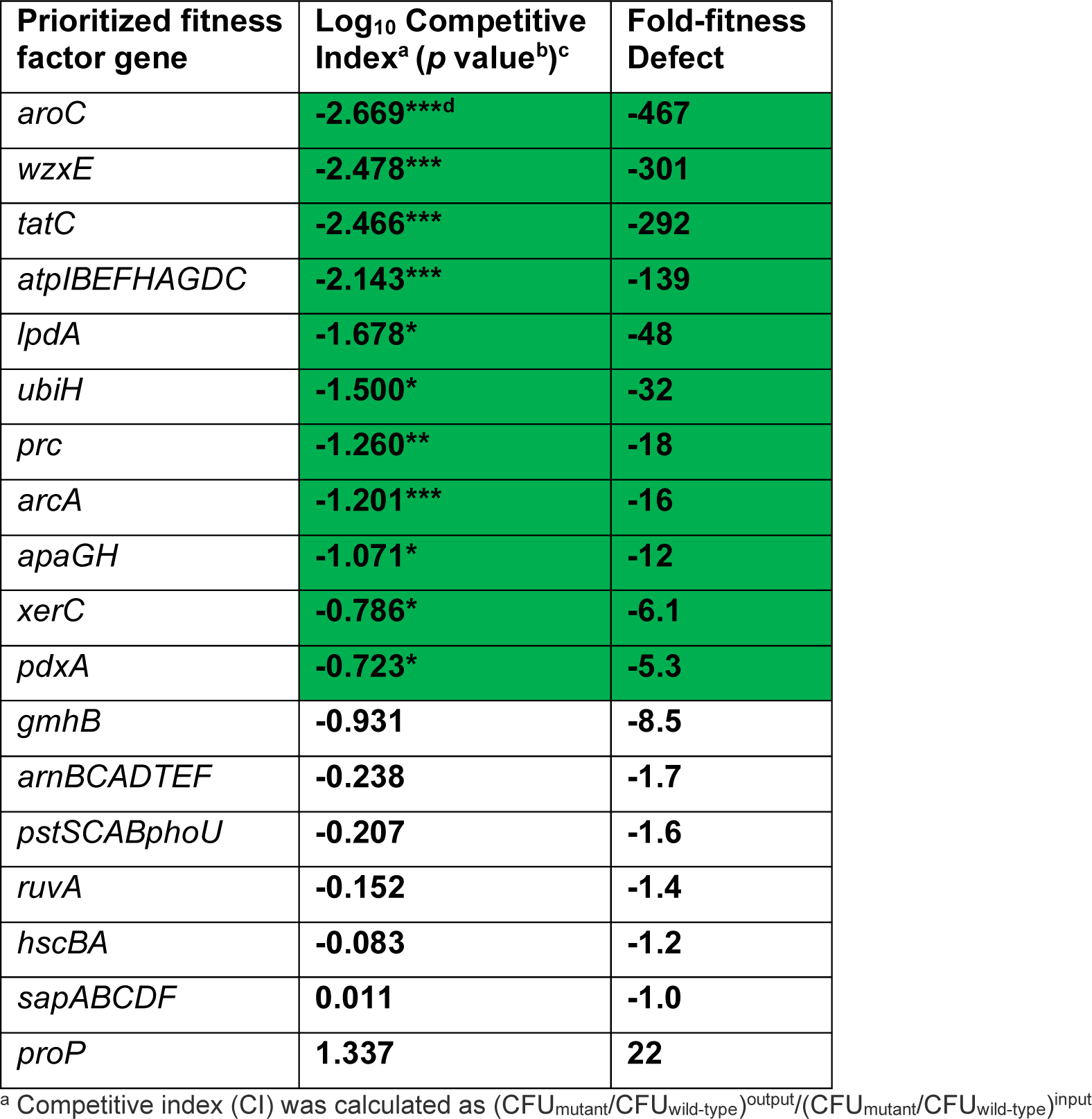
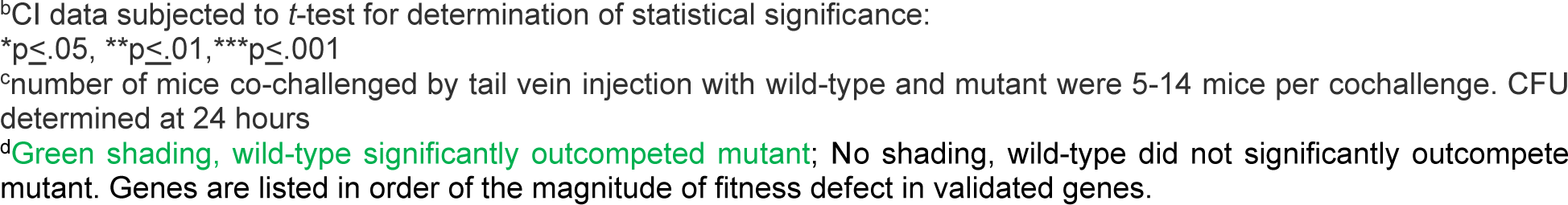
Competitive indices of prioritized Tn-Seq fitness factor gene mutants in the <u>Kidneys</u> following cochallenge with wild-type *Serratia marcescens* UMH9 in the murine model of bacteremia.

| Prioritized fitness factor gene | Log<sub>10</sub> Competitive Index<sup>a</sup> (*p* value<sup>b</sup>)<sup>c</sup> | Fold-fitness Defect |
| --- | --- | --- |
| *aroC* | -2.669****<sup>d</sup> | -467 |
| *wzxE* | -2.478*** | -301 |
| *tatC* | -2.466*** | -292 |
| *atpIBEFHAGDC* | -2.143*** | -139 |
| *lpdA* | -1.678* | -48 |
| *ubiH* | -1.500* | -32 |
| *prc* | -1.260** | -18 |
| *arcA* | -1.201*** | -16 |
| *apaGH* | -1.071* | -12 |
| *xerC* | -0.786* | -6.1 |
| *pdxA* | -0.723* | -5.3 |
| *gmhB* | -0.931 | -8.5 |
| *arnBCADTEF* | -0.238 | -1.7 |
| *pstSCABphoU* | -0.207 | -1.6 |
| *ruvA* | -0.152 | -1.4 |
| *hscBA* | -0.083 | -1.2 |
| *sapABCDF* | 0.011 | -1.0 |
| *proP* | 1.337 | 22 |

<sup>a</sup> Competitive index (CI) was calculated as $(\text{CFU}_{\text{mutant}}/\text{CFU}_{\text{wild-type}})^{\text{output}}/(\text{CFU}_{\text{mutant}}/\text{CFU}_{\text{wild-type}})^{\text{input}}$
<sup>b</sup>CI data subjected to *t*-test for determination of statistical significance:
\**p*≤.05, \*\**p*≤.01, \*\*\**p*≤.001
<sup>c</sup>number of mice co-challenged by tail vein injection with wild-type and mutant were 5-14 mice per cochallenge. CFU determined at 24 hours
<sup>d</sup>Green shading, wild-type significantly outcompeted mutant; No shading, wild-type did not significantly outcompete mutant. Genes are listed in order of the magnitude of fitness defect in validated genes.

In the spleen (**Table 3**), four mutants (*tatC, ruvA, gmhB, xerC*) were attenuated in all five species. Four mutants (*wzxE, arcA, prc, apaGH*) were attenuated in four species. Six mutants (*atpIBEFHAGDC, lpdA, ubiH, aroC, pdxA, hscBA, pstSCABphoU*) were outcompeted by wild-type in two or three bacterial species. One mutant was outcompeted in only one species (*sapABCDF*). Two mutants (*arnBCADTEF*, *proP*) were not attenuated in any species.

In the liver (**Table 4**), four mutants (*tatC, ruvA, gmhB, aroC*) were outcompeted by the wild-type of all five bacterial species. Four mutants (*ubiH, xerC, arcA, wzxE*) were attenuated in four species. Three mutants (*apaGH, hscBA, prc)* were outcompeted by wild-type in three bacterial species. Five mutants (*pstSCABphoU, lpdA, arnBCADTEF, atpIBEFHAGDC, pdxA*) were attenuated in 1 or 2 species. As for the liver, only two mutants (*proP, sapABCDF*) were not attenuated for any species.

Unlike the other four species, *S. marcescens* has an affinity for kidney colonization in addition to the spleen and liver following challenge by tail vein injection of mice (6). Eleven of the 18 mutants (*aroC, wzxE, tatC, atpIBEFHAGDC, lpdA, ubiH, prc, arcA, apaGH, xerC*, and *pdxA*) were all significantly outcompeted by the *S. marcescens* wild-type strain in the kidneys with fitness defects ranging from 5.3-fold to 467-fold (**Table 5**).

### Establishing phenotypes of mutants *in vitro*

Bacterial constructs with mutations in single prioritized fitness genes or operons across all species were then evaluated for *in vitro* phenotypes. (**Table 2, Supplemental Table 2**). We also sought *in vitro* correlates that could predict successful colonization by a bacterial species in the murine model of bacteremia. We attempted to determine whether the functional pathways identified represent a global strategy of Gram-negative bacilli to successfully infect the bloodstream.

i. **Growth rates of mutants.** Growth rates of wild-type strains and mutants were assessed during exponential growth in Luria broth at 37°C with aeration to identify potential growth defects in rich medium displayed by the mutants as compared to their respective wild-type strains. Growth was monitored over 16 h in automated growth curve analyzers. Relative growth rates were defined during exponential growth phase as maximal specific growth rate of mutant / maximal specific growth rate of wild-type (**Table 6) (Supplemental Fig 1)**. For 65 of 89 (73%) mutants, no statistically significant defect in growth rate was noted as compared to the wild-type strain. Among the strains with growth defects, mutations in a component of the pyruvate dehydrogenase complex (*lpdA*) and an enzyme in the ubiquinone synthesis pathway (*ubiH*) conferred significantly slower growth in all five species. Mutations in genes encoding ATP synthase (*atpIBEFHAGDC*) and the *pstSCAB phoU* locus similarly resulted in significantly slower growth in three of the five species. Although most of the observed growth phenotypes were consistent between species, *wzxE* mutations were an exception, resulting in enhanced growth for *S. marcescens* but slower replication for *E. hormaechei*.
ii. **ii. Susceptibility to Ciprofloxacin.** Mutants in recombinases encoded by *ruvA* and *xerC*, responsible for DNA repair, have been shown to be susceptible to the action of the DNA topoisomerase and gyrase inhibitors including the commonly prescribed antibiotic ciprofloxacin (18). We measured the diameters of zones of inhibition (killing) on agar plates around paper disks saturated with 5 µg ciprofloxacin. *ruvA* mutants of all five species and *xerC* mutants of four species had a significantly larger zone size (p<.05) to ciprofloxacin than the wild-type strains (**Table 7**). Notably, *E. hormaechei* UM_CRE_14 is completely resistant to ciprofloxacin and while mutation of *xerC* did not alter that intrinsic resistance, loss of *ruvA* did result in a significant, albeit modest, increase in susceptibility. This confirms the expected *in vitro* phenotype of these mutants and their prioritization as potential fitness factors that also affect antibiotic susceptibility.
iii. **iii. Serum Resistance.** We hypothesized that three fitness factor genes or operons may contribute to resistance to human serum. The *wzxE* mutants are expected to have reduced amounts of Enterobacterial Common Antigen (ECA) on the surface that may weaken outer membrane integrity and modify interaction with serum components including complement. *prc* encodes a protease involved in the regulation of peptidoglycan synthesis (19) but has also been demonstrated to degrade complement (20). The *sap* locus encodes proteins that provide “<u>s</u>ensitivity to <u>a</u>ntimicrobial <u>p</u>eptides” as shown in *Salmonella* Typhimurium (21). To test for serum sensitivity, 10^7^ cfu/ml of the wild-type strains and their respective *wzxE*, *prc*, and *sapABCDF* mutants were incubated with human serum for 90 minutes at 37°C and then plated for viability on Luria agar. *prc* deletion mutants of four species, excepting *K. pneumoniae* KPPR1, were found to be more sensitive to active human serum as compared to the wild-type strains (**Fig 2**). *wzxE* mutants in all five species were more susceptible to serum killing than their wild-type strains. None of the *proP* mutants were more sensitive than respective wild-type strains for any species tested. Mutants for all species were not sensitive to heat-inactivated human serum.
iv. **Susceptibility to Antimicrobial Peptides.** We tested *wzxE*, *arn*, and *sap* deletion mutants and their respective wild-type strains for susceptibility to the model cationic antimicrobial peptide polymyxin B (**Fig 3**). *arnBCADTEF* deletion mutants were statistically more susceptible to polymyxin B in all species tested (*E. hormaechei* UM_CRE_14 lacks the *arn* operon and was not tested), as were *wzxE* mutants in four species (*E. coli* CFT073, *C. freundii* UMH14, *S. marcescens* UMH9 and *E. hormaechei* UMCRE14). Unlike the published report for *Salmonella* Typhimurium (21), and opposite to the anticipated result, *sap* operon deletion mutants were statistically protected against polymyxin B relative to their respective wild-type strain in four species (*E. coli* CFT073, *K. pneumoniae* KPPR1, *S. marcescens* UMH9 and *C. freundii* UMH14). Finally, neither the *sap* operon nor the single *sapC* locus mutants in *E. hormaechei* UM_CRE-14 were statistically different than the wild-type strain.
v. **Secretion of folded proteins.** SufI is a model protein substrate of the twin-arginine translocation (Tat) protein export system whose relationship with Tat secretion has been studied extensively in *E. coli* (22-24). The Tat pathway is widely conserved among bacteria (25) and we have previously demonstrated that both *tatC*, encoding twin-arginine signal peptide recognition capacity, and *sufI* were significant contributors to bacteremia fitness of *C. freundii* (11). Thus, we chose to demonstrate its phenotype in an uninvestigated species. In wild-type exponentially growing *S. marcescens* expressing a SufI-GFP fusion, a strong fluorescent signal was observed at one or both cell poles (**Fig 4A**) by fluorescence microscopy, resulting in increased signal intensity at these locations when plotted as a function of cell length (**Fig 4B**). This polar localization of SufI-GFP was completely *tatC*-dependent since fluorescence resulting from the same fusion construct was uniformly diffuse throughout the cell in the UMH9 Δ*tatC*::*km* strain. Similarly, differential localization was also not observed in wild-type or mutant bacteria expressing the unmodified GFP control plasmid, indicating that the SufI N-terminal signal sequence was required for the polar localization of SufI-GFP. Together, these results are consistent with Tat-dependent translocation of *S. marcescens* SufI that is mediated by the predicted twin-arginine signal sequence. In addition to the diffuse fluorescence phenotype, the Δ*tatC*::*km* mutant also exhibited a significant increase in total cell length compared to wild-type bacteria (**Fig 4C**). These findings are consistent with observations of *tat* null mutations in other species and is likely a consequence of inappropriate localization of cell division-related proteins that are secreted via the Tat pathway (23, 26-28).
vi. **Repression of aerobic growth.** Mutation of *arcA* in *E. coli*, *K. pneumoniae, S. marcescens, C. freundii, and E. hormaechei* resulted in a small colony phenotype compared to each respective wild-type strain. This was quantified by plating dilutions of cultures of *arcA* mutants and wild-type strains on Luria agar and measuring colony diameters after overnight incubation at 37°C using ImageJ-2 software (**Fig 5**). Wild-type colony diameters for *E. coli*, *K. pneumoniae, S. marcescens, C. freundii*, and *E. hormaechei* averaged 2.05 mM, 2.53 mM, 1.54 mM, 1.61 mM, and 1.56 mM, respectively, whereas *arcA* mutant colony diameters averaged 1.00 mM, 1.41 mM, 0.76 mM, 1.02 mM, and 0.72 mM (*p*<.0001 for all comparisons), respectively.
vii. **Siderophore production**. *aroC* encodes chorismate synthase which provides the precursor for the synthesis of catechol siderophores, including enterobactin (29). Thus, mutation of *aroC* prevents enterobactin biosynthesis. While *C. freundii* UMH14 synthesizes enterobactin as its only siderophore, the other four species carry multiple siderophore biosynthesis pathways. We examined total siderophore activity on CAS agar plates for wild-type strains and *aroC* mutant constructs (Fig 6). In the CAS assay, a yellow halo around each colony, resulting from chelation of iron by the siderophore from the dye, is indicative of siderophore activity. For four of the five species, wild-type strains appeared to have more siderophore activity than their respective *aroC* mutants, ranging from subtle to dramatic. For example, *S. marcescens* UMH9 (30), which synthesizes chrysobactin and the catecholate siderophore serratiachelin, showed a dramatic reduction in halo radius and intensity in the mutant compared to wild-type, whereas *K. pneumoniae* KPPR1 (31), which produces enterobactin, salmochelin, and yersiniabactin (32), and *C. freundii* UMH14, which synthesizes only enterobactin [assessed by antiSmash 7.0 (32)], showed a more subtle diminishment of halo intensity. For *E. coli* CFT073 which carries at least four siderophore biosynthetic pathways including enterobactin, salmochelin, yersiniabactin, and aerobactin as well as heme receptors (33), the loss of enterobactin, salmochelin and perhaps yersiniabactin (catechol siderophores) reduced the radius of the halo around the colony on CAS agar but the bacterium still produces significant chelating activity due to remaining siderophores. *E. hormaechei* UM_CRE_14, which encodes the ability to synthesize enterobactin (100% match for biosynthesis genes by antiSmash 7.0) and perhaps aerobactin (60% match) did not display a demonstrable difference between wild-type and mutant in this assay.
viii. **Oxidative Stress assessed by exposure to H2O2.** Resistance to oxidative stress is a key mechanism used by pathogens to evade immune responses. To determine if fitness genes annotated as involved in oxidative stress responses protected against this threat, we exposed the *arcA, ruvA,* and *xerC* mutant constructs to hydrogen peroxide **(Supplemental Fig 2)**. None of these mutant constructs were more susceptible to H2O2 across any species. Previous literature demonstrated that the gene *sspA* is required for *K. pneumoniae* oxidative stress resistance (8), although this gene is not traditionally annotated as involved in this stress response (8). Oxidative stress resistance may be conveyed by other fitness genes, and each species likely has unique mechanisms to combat this response.
ix. **Osmotic stress.** Sensitivity to osmotic stress, encountered in the bloodstream, was assessed by incubating an independent suspension (10^7^ CFU/mL) of wild-type, *proP, prc*, and *wzxE* deletion mutants in PBS, pH 7.4 with and without 2 M D-sorbitol for 30 minutes at 37°C. Viable colonies were enumerated by dilution plating onto Luria agar (**Fig 7**). *prc* mutants of *S. marcescens* UMH9 and *C. freundii* UMH14 were significantly more sensitive to osmotic stress induced by 2M D-sorbitol than their wild-type counterparts. No significant sensitivity in any *proP* deletion mutant to sorbitol was observed. In addition, *wzxE* deletions of *E. coli* CFT073, *K. pneumoniae* KPPR1, *S. marcescens* UMH9, and *E. hormaechei* UM_CRE_14 were significantly more sensitive to sorbitol than their wild-type strains.
x. **Envelope stress**. We measured the growth of *ruvA, tatC, gmhB*, and *wzxE* mutants and their respective wild-type strains in four of the five species (*C. freundii, E. coli, S. marcescens*, and *K. pneumoniae*) on MacConkey agar, a medium containing bile salts, and compared survival to growth on Luria agar by CFU enumeration (**Fig 8**). Except for *C. freundii*, wild-type strains were resistant to bile salts, which may be encountered in the liver during infection. Mutation of *gmhB* resulted in a profound sensitivity to bile salts in *E. coli*. *ruvA* mutants of *E. coli* and *K. pneumoniae* were both intermediately sensitive. For *tatC* mutants, *K. pneumoniae* was most sensitive and for *wzxE,* both *C. freundii* and *S. marcescens* were nearly completely susceptible, with scarce CFU encountered on MacConkey agar.
xi. **Phosphate transport**. Mutation of the primary high affinity phosphate ABC transport system (*pstSCABphoU*) results in limitation of phosphate, a requisite nutrient (34). When bacteria experience phosphate starvation, they upregulate alkaline phosphatase encoded by *phoA*, which scavenges phosphate from monoesters of phosphoric acid (35). Thus, *pst* mutants for all species and respective wild-type strains were assayed for alkaline phosphatase activity following culture in phosphate-limiting minimal medium (36). In sharp contrast to their wild-type strains, *pstSCABphoU* mutants of *E. coli* CFT073, *K. pneumoniae* KPPR1, *C. freundii* UMH14, *S. marcescens* UMH9 but somewhat less so for *E. hormaechei* UM_CRE_14, displayed significantly higher alkaline phosphatase activity when cultured in this phosphate limited condition (**Fig 9**).

**Fig 2.**
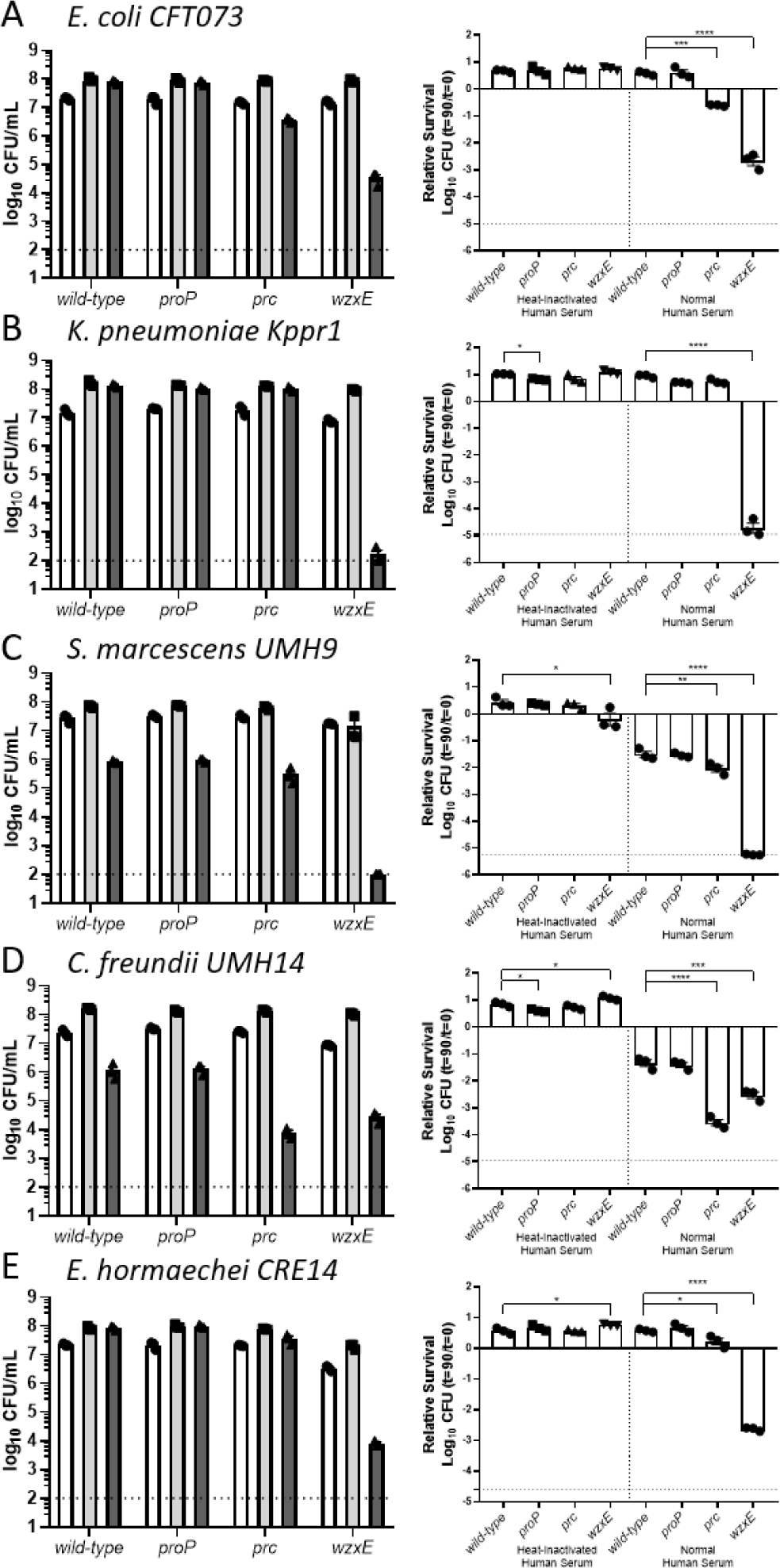
Susceptibility of Enterobacterales species to human serum. 1×10^7^ CFU/mL of bacteria (A-E: named species and strain) were incubated with normal human serum and heat-inactivated human serum to indicate complement-specific killing for 90 minutes at 37°C. 90% pooled human serum was used for *K. pneumoniae* KPPR1 and 40% pooled human serum for *C. freundii* UMH14*, E. coli* CT073, *S. marcescens* UMH9, and *E. hormaechei* UM_CRE_*14*. (Left panels) Individual CFUs with mean +/− SEM (n=3) were plotted at t=0 (white) and t=90 in heat-inactivated human serum (light gray) and normal human serum (dark gray). (Right panels) Viability was calculated relative to t=0 with statistical differences in susceptibility to heat-inactivated human serum or normal human serum determined using an unpaired *t*-test (\**p*<0.05, \*\**p*<0.01, \*\*\**p*<0.001, \*\*\*\**p*<0.0001). Dashed line denotes limit of detection. Data are presented as the mean ± SEM and are representative of 3 independent experiments each with 3 biological replicates.

**Fig 3.**
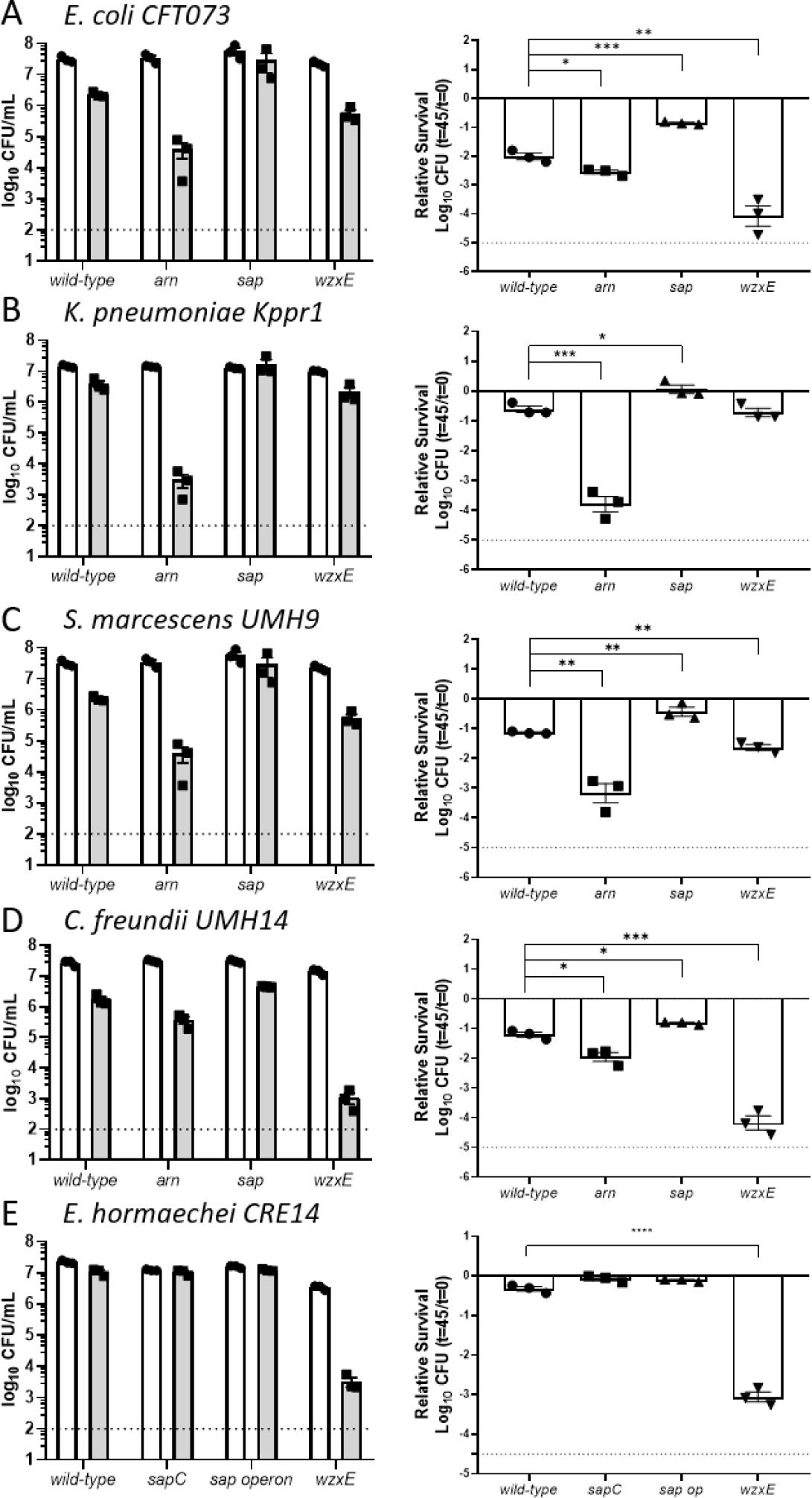
Susceptibility of Enterobacterales species to Polymyxin B. 1×10^7^ CFU/mL of bacteria (A-E: named species and strain) were incubated with Polymyxin B in PBS for 45 minutes at 37°C. (Left panels) Individual CFUs with mean +/− SEM (n=3) were plotted at t=0 (white) and t=45 (gray). (Right panels) Viability was calculated relative to t=0 with statistical differences in sensitivity to Polymyxin B determined using an unpaired *t*-test (\**p*<0.05, \*\**p*<0.01, \*\*\**p*<0.001, \*\*\*\**p*<0.0001). Polymyxin B concentrations used are as follows: 1 µg/mL for *E. coli* CFT073; 2.5 µg/mL for *C. freundii* UMH14; 5 µg/mL for *K. pneumoniae* Kppr1; 10 µg/mL for *S. marcescens* UMH9 and 1 µg/mL for *E. hormaechei* UM_CRE_14. *sap* operon indicates *sapBCADTEF* mutant. *arn* indicates *arnABCDF* mutant. Dashed line denotes Limit of Detection. Data are presented as the mean ± SEM and are representative of 3 independent experiments each with 3 biological replicates. Statistical significance was assessed by the *t*-test.

**Fig 4.**
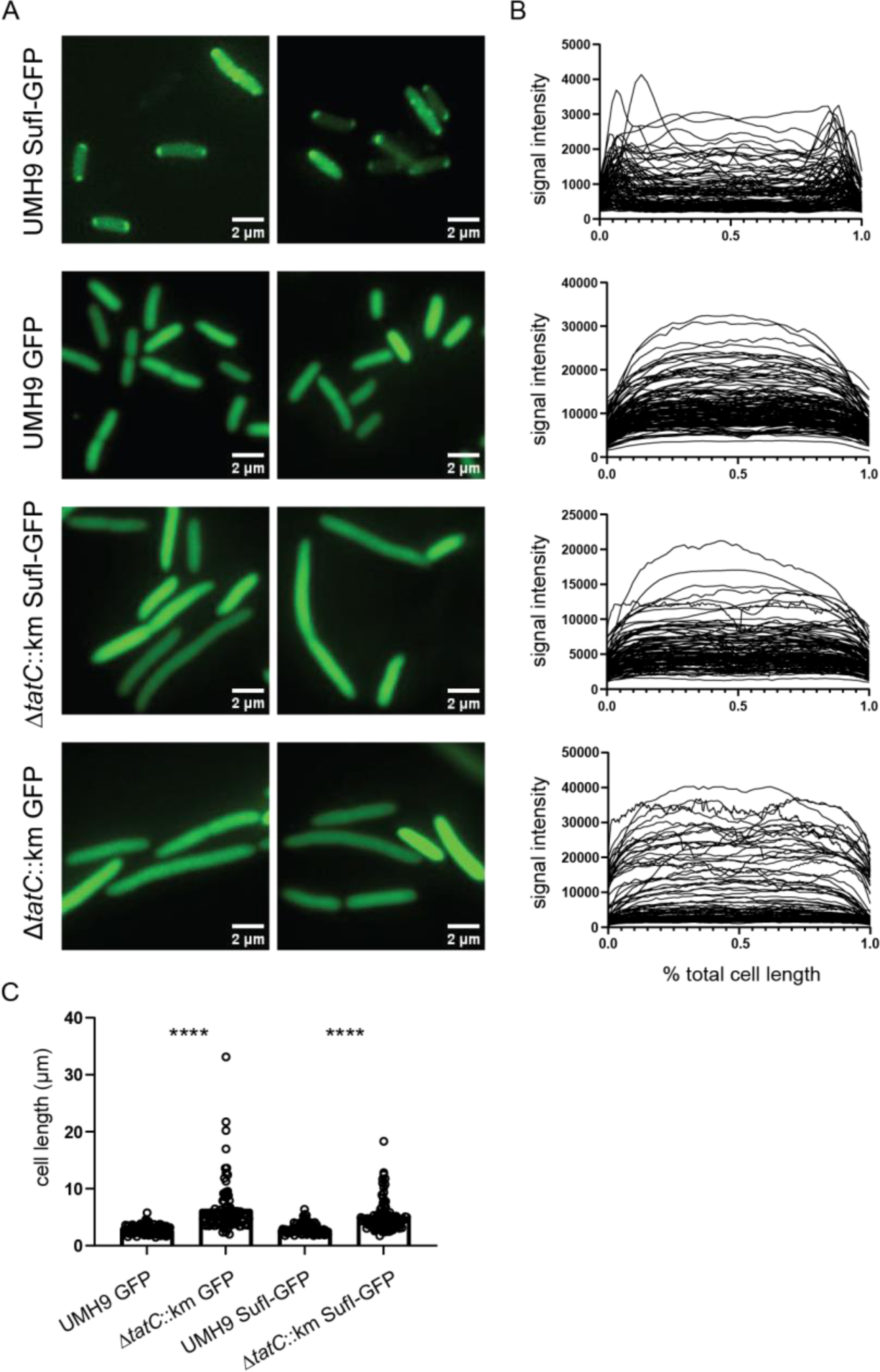
Demonstration of Tat pathway secretion in *S. marcescens*. A. *S. marcescens* wild-type and *tatC* mutant bacteria harboring plasmids that encoded an unmodified GFP (GFP) or an engineered fusion of the SufI N-terminal twin-arginine signal peptide with GFP (SufI-GFP) were visualized by fluorescence microscopy. Fluorescence intensity as a function of cell length (B) and total cell length (C) were determined for bacteria (n≥100) from multiple fields using Image J. Statistical significance for panel C was assessed by unpaired *t*-test: ****, *p*<0.0001.

**Fig 5.**
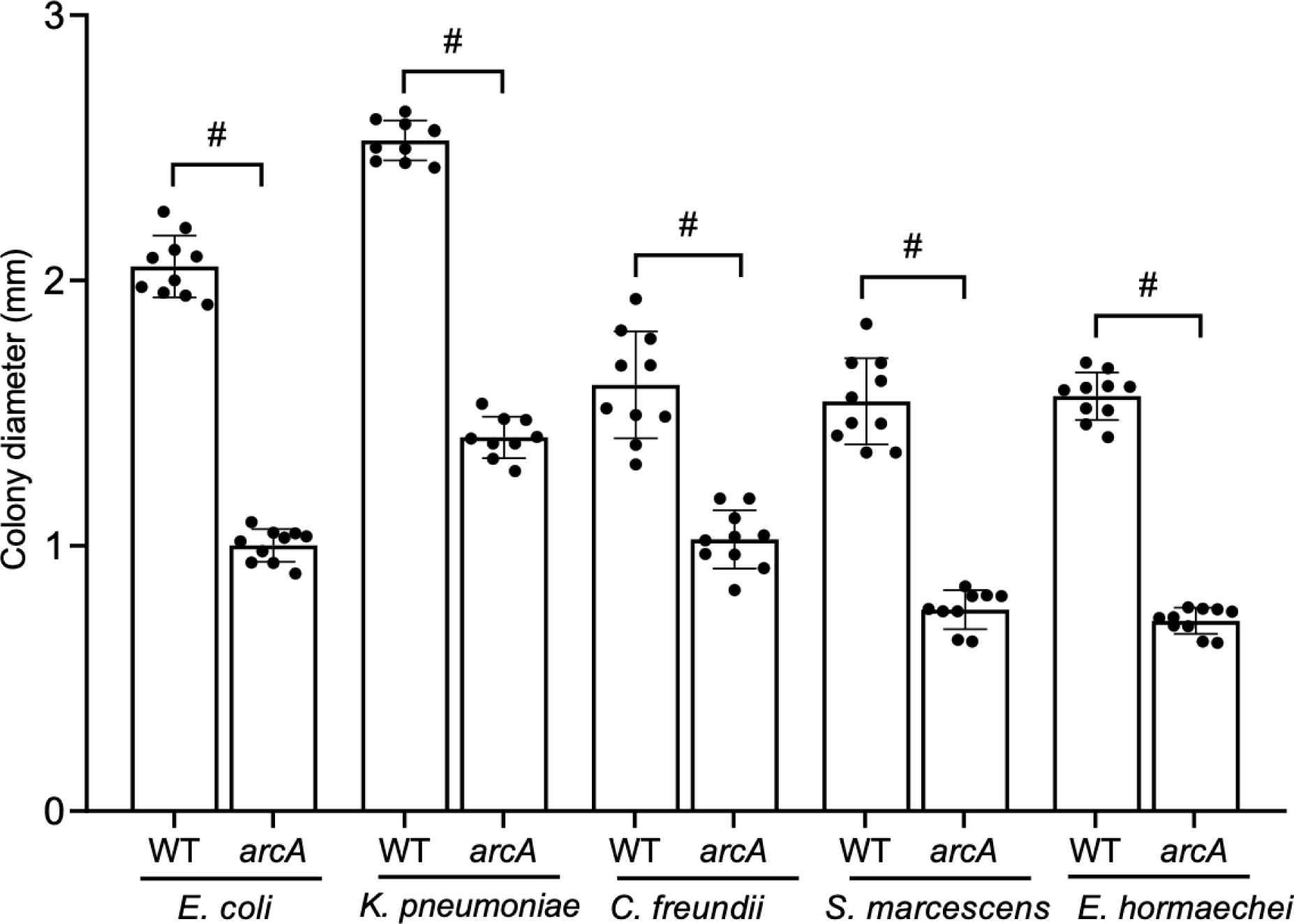
Small colony phenotype of *arcA* mutants. Wild-type and *arcA* mutants were cultured5overnight in Luria broth with aeration at 37°C. Ten-fold dilutions were spread plated onto Luria agar and incubated overnight at 37°C. Colony diameters were measured using ImageJ software (http://imagej.nih.gov/ij). ^#^For all five species, *arcA* mutants had statistically significantly smaller colony diameters than wild-type strains as determined using an unpaired *t*-test. ^#^(p<.0001).

**Fig 6.**
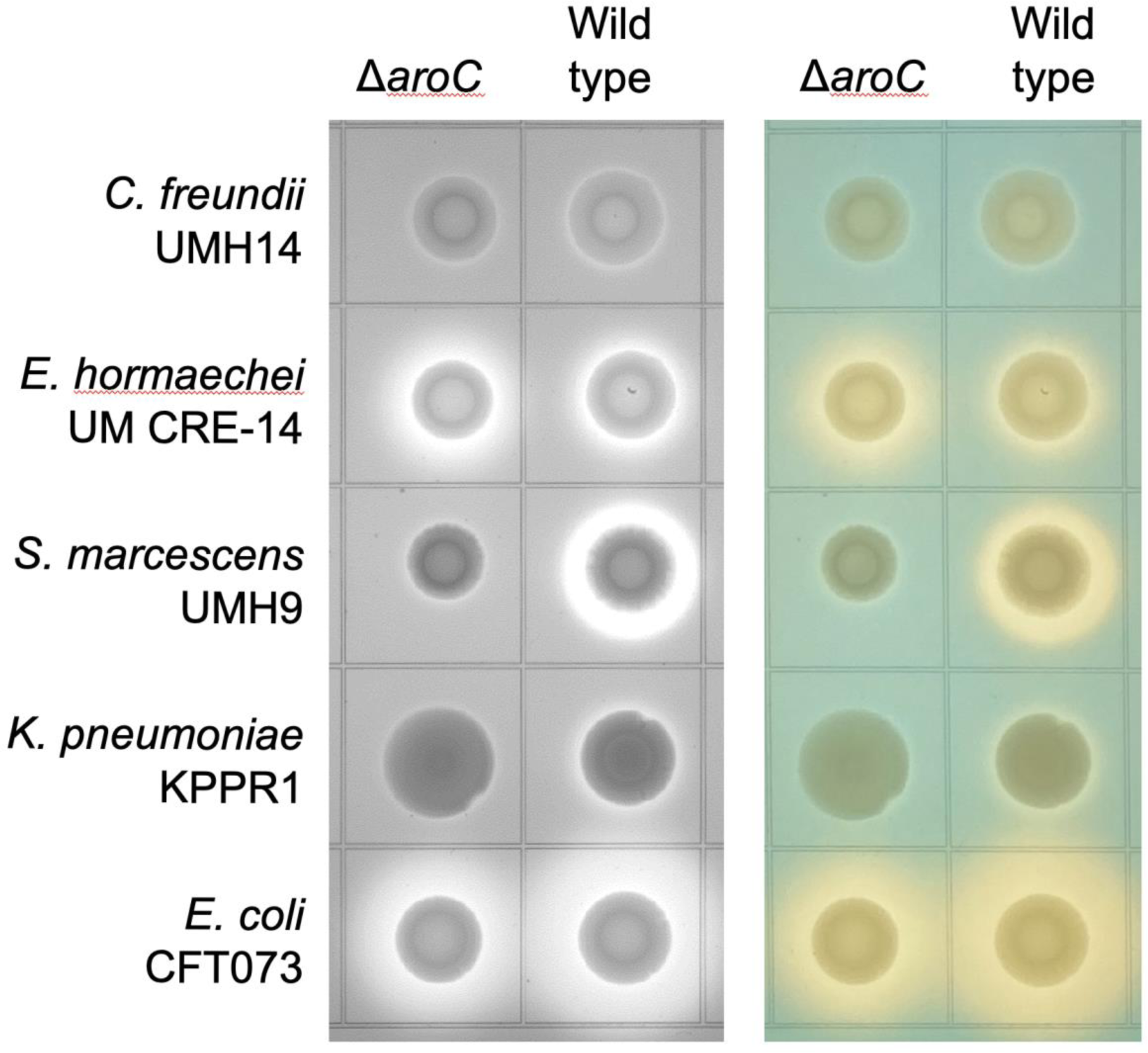
Siderophore production. Siderophore production was detected on chrome azurol S (CAS) plates supplemented with 1% tryptone. A sample (2 µl) of overnight stationary phase LB medium cultures was spotted on to CAS agar plates and incubated at 37°C for 16 h. Siderophore activity was indicated by a shift from blue to yellow color surrounding colonies. Both black and white and color images of the same plate are included to detect subtle difference in the sharpness of zones beyond the colonies and the intensity of the color change.

**Fig 7.**
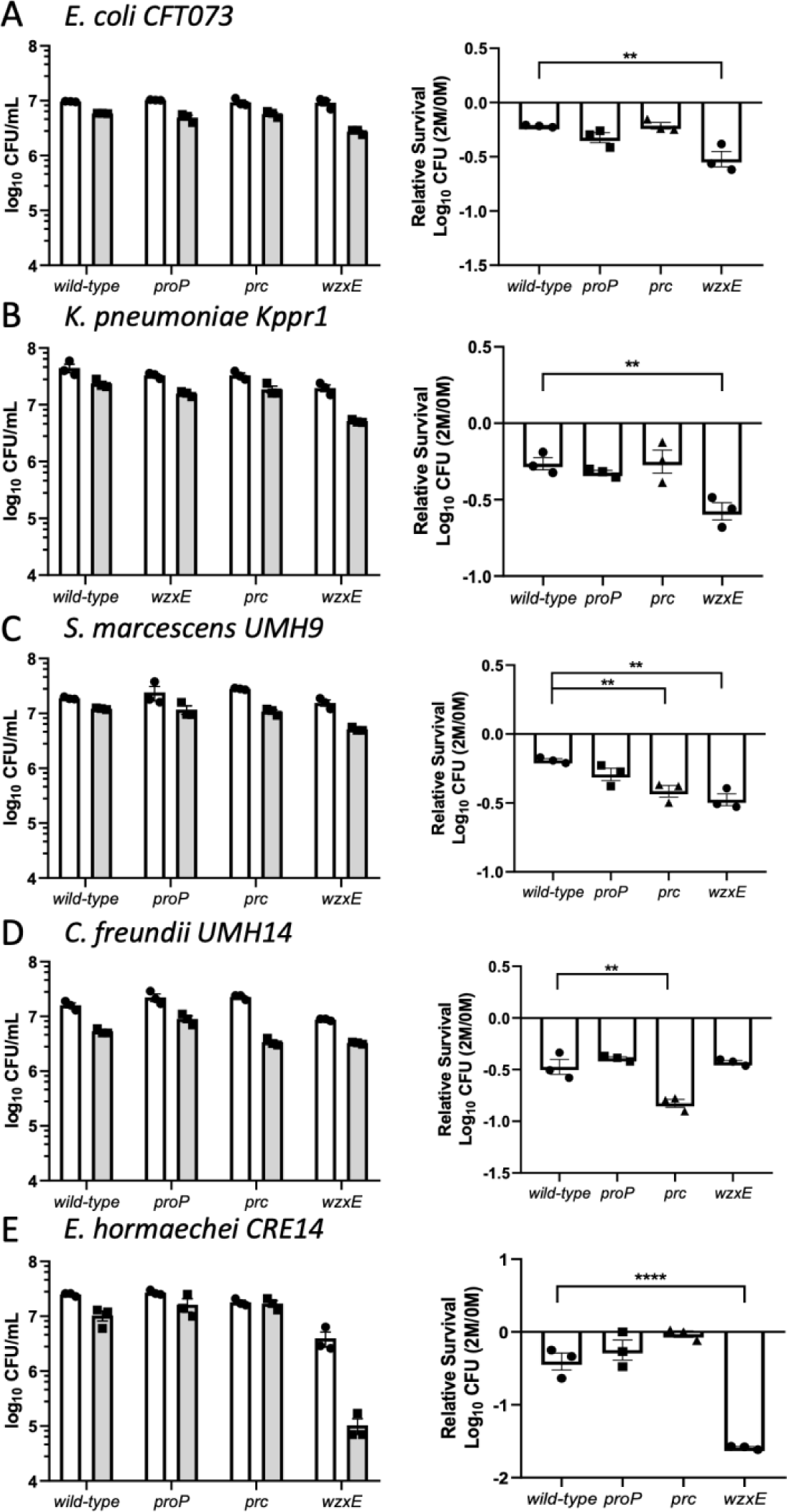
Susceptibility of Enterobacterales species to osmotic stress. 1×10^7^ CFU/mL of bacteria (A-E: named species and strain) were incubated with 0M or 2M D-sorbitol in PBS to induce osmotic stress for 30 minutes. (Left panels) Individual CFUs with mean +/− SEM (n=3) were plotted after 30-minute incubation in 0M (white) and 2M (gray) sorbitol. (Right panels) Bacterial viability was calculated relative to 0M sorbitol. Data are presented as the mean ± SEM and are representative of 3 independent experiments each with 3 biological replicates. Statistical significance was assessed by an unpaired *t*-test.

**Fig 8.**
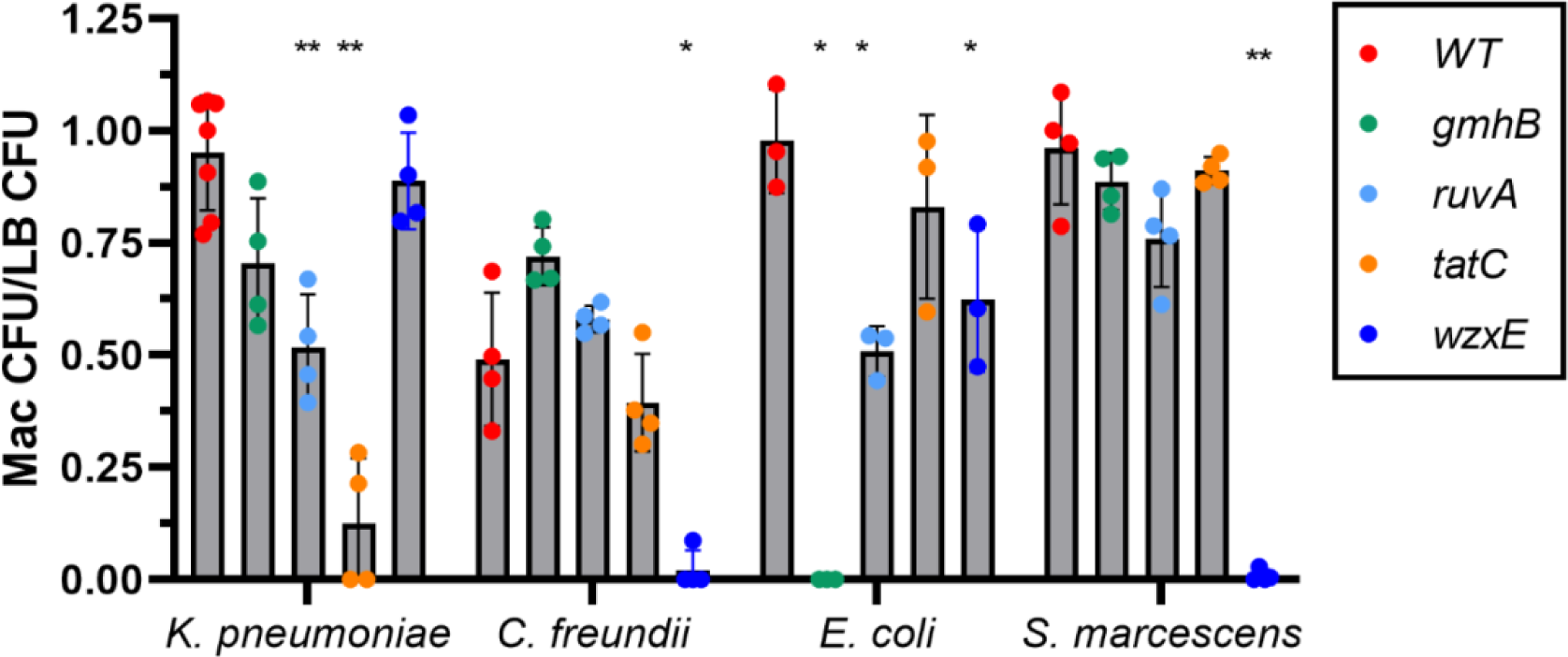
Envelope stress after exposure to bile salts. Bacterial cultures were incubated at 37℃ overnight in Luria broth, diluted in PBS to a final concentration of 10^4^ CFU/mL. Bacterial suspensions were spread-plated onto MacConkey agar (Mac) and LB agar (LB) in triplicate and incubated overnight at 27°C Colonies were counted after 48 hours of incubation. Data are presented as the ratio of the number of CFU on MacConkey agar to the number of CFU on LB agar (78). Significance was determined by paired ANOVA Dunnett’s multiple comparisons test. *, *p*<.05; **, *p*<.01.

**Fig 9.**
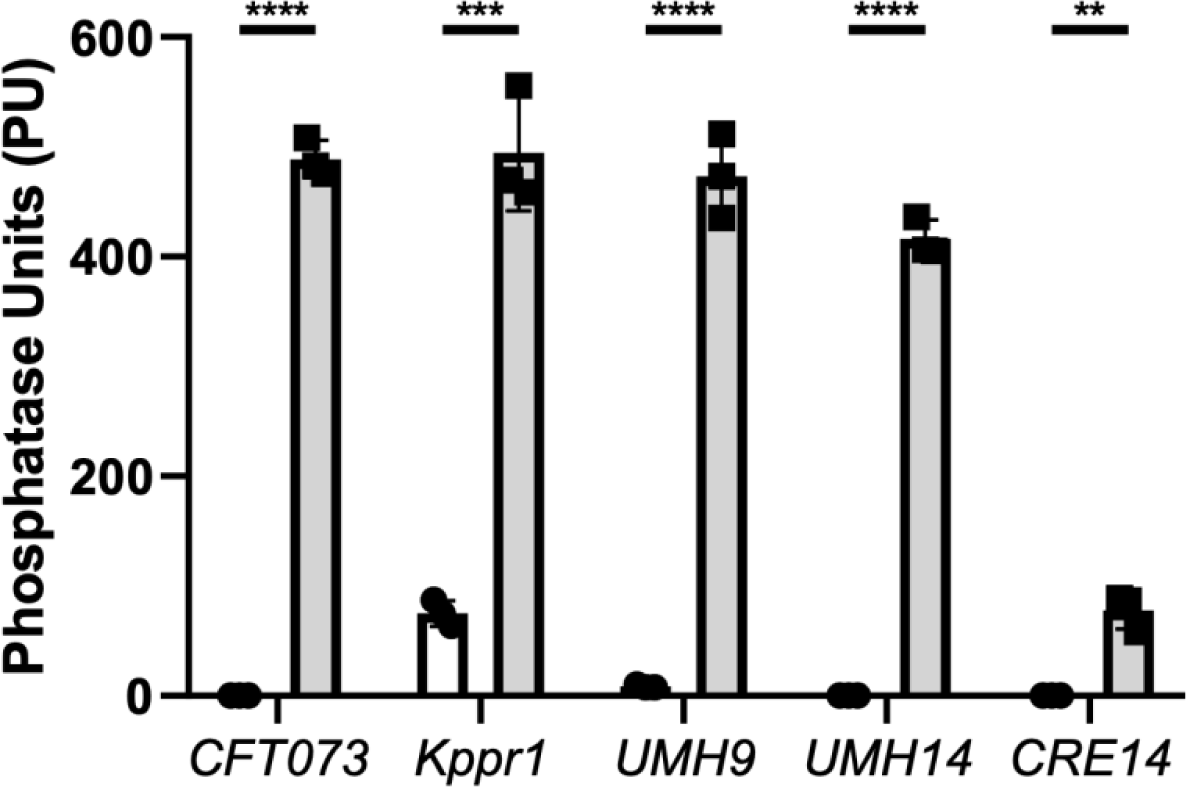
Alkaline phosphatase activity of phosphate transport mutants. Suspensions of wild-type strains normalized to OD_600_=0.1 (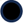) *E. coli* CFT073, *K. pneumoniae* Kppr1, *S. marcescens* UMH9, *C. freundii* UMH14, *E. hormaechei* UM_CRE_14 and their respective *pstSCABphoU* mutants (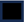) were incubated with *p*-nitrophenylphosphate at 37°C for one hour. Hydrolysis of substrate was followed OD_405_ as a measure of alkaline phosphatase activity. Statistical differences were measured with an unpaired *t*-test and (\*\**p*<0.01, \*\*\**p*<0.001, ****I<0.0001).

**Table 6.** Relative growth rates of mutants compared to wild-type strains in LB medium^a^.

| Mutant | Mean Relative Growth Rate<sup>b</sup> (Adj. P<sup>c</sup>) |  |  |  |  |
| --- | --- | --- | --- | --- | --- |
|  | *E. coli*<br>CFT073 | *K. pneumoniae*<br>KPPR1 | *S. marcescens*<br>UMH9 | *C. freundii*<br>UMH14 | *E. hormaechei*<br>UM_CRE_14 |
| *apaGH* | 0.88 (NS) | 1.06 (NS) | 1.06 (NS) | 0.98 (NS) | 0.98 (NS) |
| *arcA* | (16)<sup>d</sup> | (16) | 0.79 (NS) | (16) | 0.51 (<0.0001) |
| *arnBCADTEF* | 0.89 (NS) | 1.09 (NS) | 1.03 (NS) | 1.09 (NS) | ND<sup>e</sup> |
| *aroC* | 0.92 (NS) | 0.66 (0.0056) | 1.18 (NS) | 0.99 (NS) | 0.96 (NS) |
| *atpIBEFHAGDC* | 0.68 (NS) | 0.74 (0.0486) | 0.62 (0.0002) | 0.87 (NS) | 0.75 (<0.0001) |
| *gmhB* | 0.67 (NS) | (13) | 1.16 (NS) | 0.78 (NS) | 0.89 (0.0016) |
| *hscBA* | 0.70 (NS) | 0.89 (NS) | 0.88 (NS) | 1.08 (NS) | 0.90 (0.0073) |
| *lpdA* | 0.25 (<0.0001) | 0.47 (<0.0001) | 0.50 (<0.0001) | 0.46 (0.0107) | 0.31 (<0.0001) |
| *pdxA* | 0.81 (NS) | (8) | 0.97 (NS) | 1.17 (NS) | 0.94 (NS) |
| *prc* | 0.76 (NS) | 0.97 (NS) | 1.36 (0.0004) | 1.03 (NS) | 0.99 (NS) |
| *proP* | 0.87 (NS) | 0.97 (NS) | 1.07 (NS) | 0.96 (NS) | 1.01 (NS) |
| *pstSCABphoU* | 0.61 (0.0382) | 0.73 (0.0346) | 0.78 (NS) | 0.73 (NS) | 0.90 (0.0106) |
| *ruvA* | 0.81 (NS) | 1.01 (NS) | 0.86 (NS) | (11) | 0.97 (NS) |
| *sapABCD* | 0.70 (NS) | 0.83 (NS) | 1.20 (NS) | 1.02 (NS) | 0.79 (<0.0001) |
| *tatC* | 0.85 (NS) | 1.04 (NS) | 0.89 (NS) | (11) | 0.95 (NS) |
| *ubiH* | 0.28 (<0.0001) | 0.36 (<0.0001) | 0.64 (0.0004) | 0.43 (0.0065) | 0.39 (<0.0001) |
| *wzxE* | 0.84 (NS) | 1.04 (NS) | 1.60 (<0.0001) | 0.90 (NS) | 0.80 (<0.0001) |
| *xerC* | 1.01 (NS) | 0.97 (NS) | 0.93 (NS) | 1.07 (NS) | 0.94 (NS) |

<sup>a</sup>aerobic growth in Luria broth at 37°C
<sup>b</sup>maximal specific growth rate was determined using AMiGA software (17). Relative growth rates (mutant/wild-type) <1 indicate mutant grew more slowly than wild-type.
<sup>c</sup>statistical significance was assessed by one-way ANOVA with Dunnett's multiple comparisons test against a reference value of 1 representing wild-type. NS, not significant.
<sup>d</sup>growth rates previously determined in the indicated reference from our group
<sup>e</sup>not determined, genes not present

**Table 7.** Zones of inhibition for disk diffusion of ciprofloxacin in wild-type strains and *xerC* and *ruvA* mutants.

| Bacterial strain | Wild-type/<br>mutant | Zone of<br>Inhibition<br>(mM)<sup>b</sup> | *P*-value<sup>c</sup> |
| --- | --- | --- | --- |
| *E. coli* CFT073 | wild-type | 21.8 |  |
|  | *xerC* | 27.3 | .0327 |
|  | wild-type | 22.6 |  |
|  | *ruvA* | 29.0 | .0007 |
| *K. pneumoniae* KPPR1 | wild-type | 21.7 |  |
|  | *xerC* | 25.7 | .0224 |
|  | wild-type | 22.3 |  |
|  | *ruvA* | 27.3 | .0004 |
| *S. marcescens* UMH9 | wild-type | 22.0 |  |
|  | *xerC* | 26.7 | .0002 |
|  | wild-type | 21.0 |  |
|  | *ruvA* | 26.3 | .0013 |
| *C. freundii* UMH14 | wild-type | 22.0 |  |
|  | *xerC* | 30.0 | .0023 |
|  | wild-type | 24.0 |  |
|  | *ruvA* | 28.7 | .0249 |
| *E. hormaechei* UM_CRE_14<sup>a</sup> | wild-type | 0.00 |  |
|  | *xerC* | 0.00 | NS<sup>d</sup> |
|  | wild-type | 0.00 |  |
|  | *ruvA* | 10.7 | <.0001 |

<sup>a</sup>*E. hormaechei* wild-type and *xerC* mutant were not susceptible to ciprofloxacin
<sup>b</sup>average of 3 replicates with disk containing 5 µg ciprofloxacin on agar plate
<sup>c</sup>Mean zone of inhibition compared using the unpaired *t*-test
<sup>d</sup>NS, not significant

## Discussion

Our previous series of Tn-Seq screens for five Gram-negative bacterial species (7-11) (Ottosen, in preparation) in the Enterobacterales order that cause a significant proportion of cases of bacteremia and sepsis in humans (3) allowed us to identify shared key fitness factor genes required for survival and replication in the bloodstream and blood-filtering organs in the murine model of bacteremia. Using genes found both in the multi-species core genome of at least four out of five species (Fouts *et al*., submitted) and identified in the Tn-Seq screens, we ranked individual fitness factor genes based on magnitude of fitness defects within individual species, frequency of occurrence among the five species, presence of multiple fitness factor genes within single operons, and conversion of the *E. coli* BW25113 (14, 15) to specific antibiotic susceptibilities when a fitness factor gene in an ordered library was mutated (**Table 2**). Using the top fitness factor genes among the five Enterobacterales species, we constructed identical mutations in 18 highly prioritized genes or operons in four species and 17 mutants in *E. hormaechei* for a total of 89 mutants. Each mutant was competed with its respective wild-type parent strain in the bacteremia model by tail vein injection of identical numbers of CFUs of wild-type and mutant bacteria. Based on the results of these studies, we reranked the fitness factor genes based on their respective significant impacts on survival (competitive indices and fold-defects) and number of species attenuated in the murine model of bacteremia (**Table 3** and **4**, **Supplemental Figure 3**).

Using survival in the spleen and secondarily in the liver, 55 of the 89 mutants were significantly outcompeted by their parent strain in the spleen (**Table 3**) and 52 of 89 mutants were significantly outcompeted by their parent strain in the liver (**Table 4**). 61 of 89 mutants were outcompeted by wild-type in either the spleen or liver. Finally, 43 of 89 mutants were outcompeted by wild-type in both the spleen and the liver. Comparing competitive indices determined in the cochallenges and the number of species attenuated by mutation of the genes, several fitness factor genes rose to the top of our consideration. For the spleen, these are: *tatC* (twin arginine transporter), *ruvA (*an endonuclease that resolves Holliday junctions (37)*, gmhB* (D,D-heptose 1,7-bisphosphate phosphatase involved in LPS synthesis)*, xerC* (tyrosine recombinase required for chromosomal segregation at cell division)*, wzxE* (flippase required for Enterobacterial common antigen synthesis)*, arcA* (regulator of aerobic respiration), and *prc (*protease involved in peptidoglycan synthesis regulation and inactivation of complement (20, 38). In the liver, a similar hierarchy of fitness factors was authenticated across the board (*tatC, ruvA, gmhB,* and *xerC)*, however, both *aroC* (a member of the shikimate biosynthesis pathway) and *ubiH* (a component of the shikimate-dependent pathway for ubiquinone biosynthesis) were also prominent fitness factor genes for all five species and four species, respectively.

Measurement of *in vitro* growth rates revealed that mutation of fitness factors genes had varying influence on doubling time (**Table 6**) **(Supplemental Fig 1)**. Indeed, most (65 of 89) of the mutants displayed no significant growth defect during *in vitro* mono-culture in LB medium but their corresponding transposon mutants had been outcompeted during the course of bacteremia. On the other hand, some mutants displayed severe growth defects. For example, mutants in the *atp* operon encoding ATP synthase where the mutation causes a severe *in vitro* growth defect by limiting the synthesis of ATP from both aerobic and anaerobic respiration. As well *ubiH* and *lpdA* mutants also revealed significant growth defect. In contrast, *wzxE* and *prc* mutants displayed significantly faster doubling times than their wild-type strains in *S. marcescens*. Regardless of the *in vitro* growth rate of the mutant, in the original Tn-Seq screens, all 18 genes were identified as bonified hits comparing *in vivo* output (spleen and liver bacterial burden) to *in vitro* input (growth of the inoculum in Luria broth). It should also be noted that growth defects in LB medium may not recapitulate the host environment since the available nutrients clearly differ between the two milieus. While the three outcomes (no change, slower, or faster growth rates) are interesting, perhaps greater weight should be given to fitness gene mutants that display no significant *in vitro* growth defects but are significantly outcompeted in the mouse model. As well, emphasis could be placed on conserved genes that are not considered housekeeping genes, based on their predicted or known functions and lack of *in vitro* growth defects. Examples of these include the virulence factors *gmhB* that is important for LPS synthesis, *wzxE* that encodes a key enzyme (flippase) in ECA synthesis, and *tatC* secretion of folded proteins by the twin-arginine transporter and factors providing protection against host defenses including *ruvA*- and *xerC*-mediated genome repair, and *prc* periplasmic protease that regulates peptidoglycan synthesis but also inactivates complement.

Certain fitness factor mutants, identified in the initial Tn-Seq screens, were validated in all or multiple species. However, because of the large scope of mutant construction, a limitation of the study may be that only selected mutants were complemented. Nevertheless, as each mutant is followed up, complementation becomes part of the assessment. Prominent examples of fitness factors are discussed below in the context of the model of bacteremia pathogenesis where seven key pathways required for the pathogenesis of bacteremia were uncovered (**Fig 10**).

**Figure 10.**
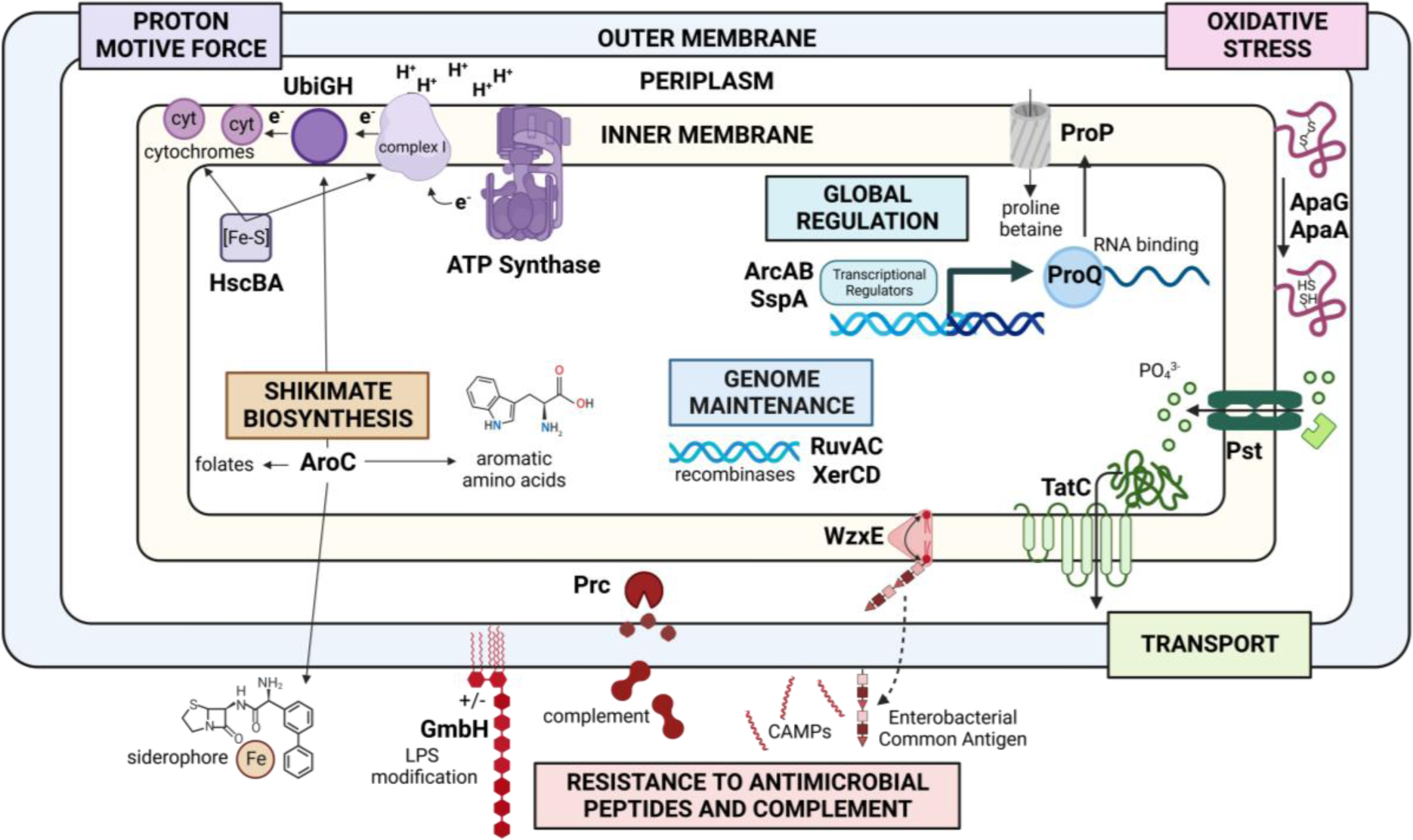
Model of Bacterial Pathogenesis of Bacteremia by Gram-negative Bacteremia. Seven common pathways are depicted that contribute to pathogenesis of bacteremia by five bacterial species within the Enterobacterales. Genes shared within the multi-species core genome of five bacterial species including *E. coli, K. pneumoniae, S. marcescens, C. freundii,* and *E. hormaechei*, were predicted as common fitness genes using Tn-Seq screens in a murine model of bacteremia. Prioritized mutants were constructed in 18 conserved genes or operons in all 5 species. Mice were cochallenged with each mutant and its respective wild-type strain by tail vein injection of mice. Genes that were validated in from 1-5 species as attenuated as measured by competitive indices are included in the model.

### Maintenance of Proton Motive Force

Mutants in ATP synthase (*atpIBEFHAGDC*) and Iron-sulfur cluster biosynthesis (*hscBA*), required for cytochrome maturation, were validated in *E. coli*, *K. pneumoniae*, and *S. marcescens* in the spleen. Ubiquinone synthesis (*ubiH*), also required for cytochrome maturation and essential for electron transport was validated in *E. coli*, *K. pneumoniae*, *S. marcescens*, and *E. hormaechei* in the liver. These data suggest that the levels of aerobic growth, generating ATP via chemiosmosis, may differ in the liver compared to the spleen. While oxygen is tightly bound by hemoglobin within erythrocytes circulating in the bloodstream, we would nevertheless consider arterial blood as a milieu that would favor aerobic growth and thus these genes would support growth in this environment.

### Resistance to Complement and Osmotic Stress

Enterobacterial Common Antigen is a carbohydrate antigen composed of repeating subunits of three amino sugars including *N*-acetylglucosamine, *N*-acetyl D-mannosaminuronic acid, and 4-acetamindo-4,6-dideoxy-D-galactose (39) that contributes to resistance to antimicrobial peptides and complement. *wzxE* mutants lacking a flippase, which translocates ECA intermediates from cytosol to periplasm, was attenuated in 4 of 5 species in the spleen and the liver, but not in *C. freundii* in either organ. The antigen is found in three forms including ECAPG that is covalently linked to phospholipids in the outer leaflet of the outer membrane envelope, ECALPS that is linked to the core polysaccharide of LPS in cases where O-antigen is not synthesized (*i.e*., in “rough” strains) and ECACYC that consists of 4-6 repeating cyclized units (depending on the species) that is localized to the periplasm (39). ECA structure is conserved in all bacterial species within the Enterobacterales order of Gram-negative non-spore forming facultatively anaerobic rod-shaped bacteria within the class *Gammaproteobacteria*. WzxE translocates the isoprenoid carrier bound to the three amino sugar precursors from the cytosol to the periplasm where it is assembled into the intact antigen and translocated to the surface of the outer membrane, anchored to the phospholipids in the case of ECAPG. ECA may represent a target of therapy as this mutant was susceptible to human serum, osmotic stress, and/or antimicrobial peptides in all species, and bile salts for three species (**Figs 2, 3, 7, and 8; Supplemental Table 2**). Although mouse serum is not bactericidal for these species, susceptibility to human serum correlated with fitness in the mouse model. This may suggest that protection from other complement-mediated defenses by *wzxE*, such as opsonophagocytosis, is important during bacteremia. In addition, *wzxE* mutants were defective in the liver and spleen and susceptible to osmotic stress in the same four species, but not in *C. freundii*. This perfect correlation suggests that *wzxE* may resist osmotic stress encountered during bacteremia. Eight other genes in the *wec* operon encoding ECA were also hit in the Tn-Seq screens (Fouts et al., submitted) highlighting the critical importance of this neglected cell surface antigen. Indeed, *wzxE* was the most highly prioritized gene and its operon was also most highly prioritized (**Table 2**) Clearly, it will be important to evaluate a whole operon deletion in the murine model of bacteremia.

The *prc* mutant was also validated as attenuated in multiple species (4 of 5 species in the spleen and 3 of 5 species in the liver). The periplasmic protease Prc has at least three roles in pathogenesis including regulation of cell wall synthesis, motility, and complement inactivation. Cross-link-specific peptidoglycan hydrolase couples the expansion of the peptidoglycan sacculus with that of phospholipid synthesis in Gram-negative bacteria. Unregulated activity of a peptidoglycan hydrolytic enzyme, MepS is detrimental for growth of *E. coli* during fatty acid-limiting conditions. Cellular availability of fatty acids or phospholipids alters post-translation stability of MepS by modulating the proteolytic activity of a periplasmic adaptor-protease complex, NlpI-Prc toward MepS (19). Mechanistically, an NlpI homodimer binds two molecules of the Prc protease and forms three-sided MepS-docking cradles using its tetratricopeptide repeats (40). However, non-metalated (apo) NDM-1 is degraded by the periplasmic protease Prc that recognizes its partially unstructured C-terminal domain. Zn(II)-binding renders NlpI is refractory to degradation by quenching the flexibility of this region (41). Prc, a bacterial periplasmic protease, and its homologues have also been directly demonstrated to be involved in the pathogenesis of Gram-negative bacterial infections. The *prc* mutant of *E. coli* RS218 exhibits a decreased ability to cause a high level of bacteremia (20) [as confirmed by us for *E. coli* CFT073, *C. freundii* UMH14, *S. marcescens* UMH9 and *E. hormaechei* UM_CRE_14 (**Tables 3 and 4**)] and is more susceptible to human serum killing than strain *E. coli* RS218 as are the aforementioned strains in this study (**Fig 2, Supplemental Table 2**). This sensitivity appears due to the mutant’s decreased ability to avoid the activation of the antibody-dependent and independent classical complement cascades as well as its decreased resistance to killing mediated by the membrane attack complex (20). Evasion of classical complement-mediated serum killing of our representative Gram-negative bacilli makes Prc a potential target for the development of therapeutic and preventive measures against Gram-negative bacteremia. Indeed, a *prc* mutant is also attenuated in experimental urinary tract infection and has reduced motility due to downregulation of FlhDC, the master regulator of flagellar synthesis (38). Thus, mutation of *prc* understandably causes multiple deleterious effects on the fitness of Gram-negative bacilli.

In this study, *gmhB* was also a prominent fitness factor gene validated in all five species although it does not fit neatly in this category as it is involved in LPS synthesis. This mutant was attenuated in all five of our Gram-negative species, being outcompeted in both the spleen and liver by their respective wild-type strains. As well, this mutant is susceptible to antimicrobial peptides in *K. pneumoniae* KPPR1 (42) and is outcompeted by wild-type during incubation in murine spleen homogenates (13). *gmbH* encodes a D,D-heptose 1,7-bisphosphate phosphatase catalyzing the third step of biosynthesis of the lipopolysaccharide (LPS) component ADP-Heptose (43, 44), essential for assembly of the LPS core on the outer membrane of Gram-negative bacilli. In a report describing the crystal structure of the enzyme, the catalytic pathway is postulated to involve a phosphoaspartate intermediate (44). It appears that GmhB is a partially redundant enzyme in the biosynthetic pathway of ADP-heptose required for the LPS core assembly. Thus, GmhB-deficient strains produce a mixed phenotype of full-length and stunted LPS molecules (44). This partial defect is attributed to an uncharacterized enzyme that is partially redundant for GmhB function. Nevertheless, some lower molecular weight components of LPS are absent in a *gmhB* mutant (13). In the bloodstream, bacterial surface structures such as intact LPS are critical to maintain outer membrane permeability and defense against complement and antimicrobial peptides (45). It is therefore curious that *gmhB* mutants of *K. pneumoniae* are not attenuated in the murine pneumonia model (13), but <u>are</u> attenuated following intraperitoneal challenge and direct inoculation into the bloodstream, either in the presence or absence of the wild-type strain. Therefore, GmhB appears to represent a promising target for therapy against bacteremia and sepsis caused by Gram-negative bacterial species as it relates to dissemination of *K. pneumoniae* and perhaps other species from sites of initial infection such as the lung or urinary tract to the bloodstream.

### Transport

The twin arginine translocation system, encoded by *tatABC*, functions in diverse bacterial species to secrete folded proteins across the cytoplasmic membrane. *tatC* mutants were attenuated in all 5 species both in the spleen and in the liver (this report and Fouts *et al*., submitted), making this transport system one the most impactful shared fitness determinants identified. The TatBC complex binds Tat substrate proteins carrying the twin arginine motif on their signal peptides (46). TatA is recruited to the activated TatBC complex and mediates transport of the substrate. TatC recognizes the secretion signal sequences for proteins that are exported through this pathway (47). We have now demonstrated that *S. marcescens* SufI subcellular localization is TatC-dependent (**Fig 4**), consistent with well-established observations in *E. coli* and other species. At least some of the fitness attenuation observed with the five *tatC* mutants in this study is likely due to this mislocalization of cell division protein SufI, as we’ve previously demonstrated for *C. freundii* (11). However, it should be noted that additional Tat-secreted proteins may also contribute to fitness and that these proteins may not necessarily be encoded in the multispecies pan-genome, since translocated proteins are expected to vary between species (47).

Phosphate, a critical nutrient, is imported by an ABC transporter encoded by the *pstSCABphoU* operon. This transporter was predicted as a potential fitness factor in 3 species. It is well known that *pst* mutants are attenuated in other species [*Proteus mirabilis* in a biofilm and UTI model (36, 48) and *Campylobacter jejuni* in the murine gut model of infection (49)]. Thus, we attempted to validate this finding for bacteremia by testing *pstSCABphoU* mutants of all five species in the murine model of bacteremia and found that two, *E. coli* CFT073 and *E. hormaechei* UM_CRE_14, were attenuated (**Fig 9**).

### Genome maintenance

*ruvA* and *xerC* DNA recombinase mutants were both outcompeted in all 5 species by each respective wild-type strain in the spleen and 4 of 5 species in the liver following cochallenge by tail vein injection (**Tables 3 and 4**). This is notable, because chromosomal DNA in bacterial cells is subjected to constant damage from exposure to physical and chemical agents (*e.g*., reactive oxygen species generated by neutrophils). Bacterial cells have developed systems to repair these defects thus preventing cell death or mutation. RuvABC enzyme complexes are involved in DNA recombination and repair and facilitate Holliday junction branch migration and resolution (50, 51). RuvABC repairs damage by catalyzing homologous exchanges between damaged and undamaged DNA (52). The protein complex resolves Holliday junction intermediates produced by RecA. RuvAB mediates branch migration of the Holliday junction and RuvC resolves the joint molecule into separate products by cleavage across the point of strand exchange (53). The protein complex is known to participate in the resolution of stalled or collapsed DNA replication forks that occur during normal chromosomal synthesis (50) though importantly, the *ruvA* mutant of *C. freundii* grew similarly to the wild-type strain during rapid replication in rich medium (11). Importantly, the *ruvA* fitness defect was confirmed by competition infection against the wild-type strain using independently constructed mutants in the five species. These mutants had no defect in survival of hydrogen peroxide exposure, although *K. pneumoniae* and *E. coli* had modest growth defects in bile salts suggesting additional stressors impinge on the Ruv system. A phenotype displayed by *ruv* mutant is the filamentation of the bacterium following UV irradiation or mitomycin C treatment (54), likely due to incomplete segregation of replicating chromosomes. Mutation of *ruvA* increased susceptibility to ciprofloxacin (**Table 7**). For *C. freundii*, transposon insertion in both *ruvA* and *ruvC* by transposon insertion resulted in a >30-fold loss of fitness (**Supplemental Figure 3**). Within the two multi-protein complexes, only RuvB was not identified as a significant fitness factor in our dataset. It is tempting to speculate that bacterial DNA damage occurring in the host environment, potentially through immune cell-mediated production of reactive oxygen species but not adequately mimicked by H2O2, may also contribute to the requirement for these complexes.

Similarly, XerC is a site-specific recombinase that resolves multimers of plasmids and also has a role in the segregation of replicated chromosomes at cell division (55). *xerC* mutants form filaments that appear unable to fully partition. XerC responds to DNA damage mediated by reactive oxygen species produced by neutrophils during infection. XerC is required for induction of the SOS response, with the result that a mutant defective in *xerC* is more susceptible to a range of DNA-damaging antibiotics including ciprofloxacin and immune-mediated killing (56). *xerC* was validated as a fitness factor in all 5 species in the spleen and 4 of 5 species in the liver (**Tables 3 and 4**). Combined, XerC and RuvA provide examples of conserved fitness factors that if disrupted by a novel inhibitor, could increase the efficacy of an FDA-approved antibiotic and sensitize bacteria to stressors in the host.

### Shikimate biosynthesis

The *aroC* mutant was attenuated in multiple species: 3 of 5 species in the spleen (**Table 3**) and 5 of 5 species in the liver (**Table 4**). The *ubiH* mutant was attenuated in 3 of 5 species in the spleen and 4 of 5 species in the liver. The *aroC* gene encodes chorismate synthase, which performs the terminal enzymatic step in the shikimate biosynthesis pathway to produce chorismate, a fundamental precursor for the biosynthesis of several biomolecules (57). In bacteria, these shikimate-dependent (*i.e*., chorismate-dependent) pathways include the biosynthesis of folates, aromatic amino acids, and the quinones, ubiquinone and menaquinone (58), which contribute to aerobic and anaerobic respiration, respectively. In addition, chorismate is also required for the biosynthesis of catecholate siderophores including enterobactin and salmochelin. In agreement with our finding that mutation of *aroC* attenuates bacterial fitness in a murine host during systemic infection, alleles for five of the seven enzymatic steps in the shikimate biosynthesis pathway, were predicted by Tn-Seq studies to contribute to bacterial fitness following TVI in the same murine model of bacteremia (Fouts *et al*.,submitted). This is also in agreement with numerous prior studies demonstrating the fundamental role of shikimate biosynthesis to bacterial pathogenesis in diverse models of disease (59-62). Fitness deficiencies resulting from mutations within the shikimate biosynthesis pathway are likely caused by the compounding effects of impaired folate biosynthesis, aromatic amino acid auxotrophy, inhibition of aerobic and anaerobic respiration, and limited iron-scavenging capacity.

The attenuation of *ubiH* mutants for growth during systemic infection illustrates the utility of aerobic respiration for energy production in the host during bacteremia. Ubiquinone plays a vital role in the electron transport chain during aerobic respiration, allowing for the utilization of oxygen as a terminal electron acceptor. (63). Chorismate is the primary substrate for the biosynthesis of ubiquinone, a nine-step enzymatic process, where UbiH (2-octaprenyl-6-methoxyphenol hydroxylase) catalyzes the sixth reaction (64, 65). Inhibition of ubiquinone biosynthesis through mutation of *ubiH* and presumably *aroC* results in reduced bacterial fitness in both spleens and livers to varying degrees across the five species tested. While all five *ubiH* mutants are profoundly reduced in their capacity for aerobic growth *in vitro*, the growth rates of *aroC* mutants are less affected. These disparate growth phenotypes suggests that alternative metabolic pathways exist to fuel ubiquinone biosynthesis in an *aroC* mutant, perhaps utilizing exogenous tyrosine in the absence of endogenous chorismate production, though this remains to be tested.

### Global Regulation

ArcA is a component of a two-component system family of bacterial transcriptional regulators, composed of sensor kinase ArcB and response regulator ArcA and senses the modulation of oxygen availability for use as an electron point during infection, undergoing fermentation during bacteremia.

In our recent work (16), *arcA* mutants were also found to exhibit a dysregulated response to changes in oxygen availability, iron limitation, and membrane perturbations, which bacterial cells may experience during infection. The genetic response of the *arcA* mutants to the cationic antimicrobial peptide polymyxin B supported an expanded role for ArcA as an activator in response to membrane damage. ArcA function was also linked to electron transport chain activity based on its response to proton motive force uncoupling by carbonylcyanide-*m*-chlorophenylhydrazone (CCCP).

### Oxidative stress

Bacteria must resist oxidative stress elicited by innate immune cells during infection. Surprisingly, no genes predicted to enhance oxidative stress resistance (*arcA, ruvA,* or *xerC*) were required for bacterial survival following *in vitro* exposure to hydrogen peroxide. However, during infection, many genes may work together to enhance oxidative stress resistance. It is likely that each species has multiple mechanisms to combat this stress during infection. For example, SspA in *K. pneumoniae* is required for resistance to oxidative stress but was not tested in these studies (8).

### Mutant not attenuated

Of the 18 bacteremia fitness loci explored in this study, only mutation of *proP* failed to produce a fitness defect in either the liver or spleen for at least one species. *proP* encodes a proline/betaine transporter for osmotic protection. While *proP per se* was not predicted in any of the Tn-Seq experiments to be a fitness factor, its regulator *proQ* was identified. Because *proQ* and *prc* are co-transcribed, we elected to explore the contribution of *proP* to bacteremia fitness to avoid potential polar effects on *prc* expression resulting from mutation of *proQ*. While *proP* mutants were not attenuated during cochallenge, we did generate a *proQ* mutant in *C. freundii* UMH14 and found it to be significantly attenuated in the bacteremia model (Log10 competitive indices in spleen and liver were −0.169 (p<.05) and −0.502 (p<.05), respectively), supporting the Tn-Seq prediction (Fouts *et al*., submitted) that *proQ* is a fitness factor.

### Model of Pathogenesis of Gram-negative bacteremia

To synthesize the substantial data set presented in this report, we can consider the following data including 1) the results of Tn-Seq screens for five Enterobacterales species surveyed in the murine model of bacteremia (**Table 1**); 2) the construction of 18 mutants for all four species (and 17 mutants for *E. hormaechei*) in genes highly predicted as attenuated in the Tn-Seq screens (**Fig 1, Table 2**); 3) competitive indices from subsequent cochallenges of wild-type and each mutant in the murine model of bacteremia (**Table 3, Table 4, Table 5**); and 4) relevant *in vitro* phenotypes for many of these mutants (**Fig 2-9**. **Tables 6** and **7**). Taken together, these data allow us to formulate a working model of the pathogenic mechanisms encoded by genes within the multi-species core genome of Gram-negative bacilli during bacteremia (**Fig 10**).

This model of pathogenesis teaches us that specific core pathways shared by Gram-negative species of the Enterobacterales are critical for infection of the mammalian bloodstream and its blood-filtering organs. First, bacteria must maintain their proton-motive force using ATP synthase to synthesize ATP using reentry of extruded protons from the periplasm into the cytosol. As well, synthesis of iron-sulfur clusters allows the maturation of cytochromes for electron transport. This implies some degree of aerobic respiration is employed by these pathogens in most cases. To that end, global regulator ArcA regulates genes associated with aerobic respiration by sensing available oxygen in conjunction with ArcB to maintain proton motive force. However, in the absence of oxygen, ArcA, with ArcB, can shift metabolism toward fermentation to survive in such an environment, a key representation of metabolic flexibility. Just as critical is the role of the outer envelope to survival in the bloodstream where bacteria must resist the toxic effects of osmotic stress and complement. The protease Prc appears to protect against complement by degrading components of the system. Enterobacterial Common Antigen (ECA) also contributes to defense against complement and antimicrobial peptides. Transport also plays a crucial role by importing phosphate through the Pst ABC transporter. As well, numerous crucial folded proteins are exported across the cytoplasmic membrane by the twin-arginine transport system in order to maintain viability in the bloodstream. Next, shikimate biosynthesis is required for three key functions that include 1) siderophore synthesis required for iron acquisition, mandatory for survival in an environment where free iron is tightly sequestered by the host; 2) quinone biosynthesis as cofactors in cytochromes for electron transport; and 3) synthesis of aromatic amino acids and folate. This pathway appears to be particularly important for survival in the liver. Bacteria must also resist oxidative stress elicited by innate immune cells including neutrophils and macrophages, which are the first line of host innate defense against infection. However, this was not directly demonstrated by resistance to H2O2 (**Supplemental Fig 2**). Finally, bacteria must maintain the integrity of their genome under stresses encountered specifically in the host compared to laboratory conditions.

Finally, we can make assertions about the metabolic flexibility of these bacterial pathogens. In studies identifying the most commonly isolated Gram-negative bacilli in cases of bacteremia, at least 80% of the species are classified as facultative anaerobes with a majority being in Order Enterobacterales while *P. aeruginosa* and *A. baumannii* are strict aerobes dominating the remainder of cases (5, 68-70). Among Gram-negative pathogens, we have shown that the temporal dynamic of bacteremia in the murine model differs by species and anatomical site (6). Despite having the same “toolbox” of major metabolic pathways at their disposal and evidently utilizing rapid replication to support infection, facultative anaerobes rely on different metabolic processes to various extents in the bacteremia model here. Revisiting why facultative anaerobes cause the largest proportion of Gram-negative bacteremia cases with this perspective poses new questions. A leading hypothesis of why they are so successful is their ability to survive in the environment and in the host focuses on metabolic adaptation (71). However, *A. baumannii* and *P. aeruginosa* are also environmental pathogens capable of successfully colonizing the bloodstream but on the other hand are strict aerobes (72, 73). Metabolic flexibility may be beneficial in that they can transition between metabolic pathways during infection, or it may perhaps refer to the use of different pathways during infection altogether. These species are possibly better described as being “metabolically advantaged”. This notion would explain why opportunistic bacteria that are facultatively anaerobic are the most common Gram-negative pathogens in cases of bacteremia, yet do not necessarily rely on the same metabolic capabilities as one another. Nevertheless, these bacteria share a common strategy to survive in the mammalian bloodstream.

## Materials and Methods

### Determination of the multi-species core genome

In preparation for the current analysis described in this report, we used a pan-genome pipeline developed by the J. Craig Venter Institute along with ∼15,000 sequenced genomes to identify the multispecies core genome shared by *E. coli, K. pneumoniae, C. freundii,* and *S. marcescens*. Using criteria of 70% amino acid sequence identity for predicted proteins, we predicted a core genome of 2850 genes shared in the multi-species core genome of these four species (Fouts, D. *et al*., submitted).

### Scoring rubric for ranking and prioritization of fitness mutants

For the fitness genes predicted by Tn-Seq in the murine bacteremia model for each of the four bacterial species listed in **Table 1** (*E. hormaechei* was excluded because Tn-Seq studies were not completed at that stage of the study), we prioritized each fitness gene for each of the four species using the Tn-Seq data, according to a scoring rubric (Fouts, D. *et al*., submitted) based on four major additive criteria: 1) the magnitude of the fitness defect associated with a gene in any one species: 3 points for fitness genes in each species’ top 20 genes; 2 points for genes ranked 21-40; and 1 point for each gene ranked 41-60; 2) a gene was a fitness factor in multiple species: 3 points for a fitness gene found in 3 or more species, 2 points if found in 2 species; 3) whether multiple fitness genes reside in the same operon: 3 points if 5-9 other fitness genes are encoded in the same operon, 2 points if 3-4 other fitness genes are encoded in the same operon, and 1 point if 1-2 other fitness factors are encoded in the same operon. An additional 1 point was awarded to fitness genes encoded in operons where >49% of the operon loci were predicted to be fitness factors; 4) mutation of a fitness gene was found to confer increased antibiotic susceptibility in *E. coli* BW25113 (15); 2 points for such a fitness gene. The sum of the four criteria above was used to assign individual scores for all fitness genes or found in each species pan-genome core. Operon scores were calculated by summing all the individual scores of fitness genes encode within that operon.

### Construction of mutants in prioritized fitness genes

Fitness gene mutations in all species were generated by lambda red recombineering using established protocols (10, 11, 59, 74). For *C. freundii*, *E. coli*, *K. pneumoniae,* and *S. marcescens* the *nptII* kanamycin resistance cassette was PCR-amplified from pKD4 (74, 75) and directed to in-frame deletions of target genes via 5′-end homologous sequences (**Supplemental Table 1**). At least six codons on the 3′-end of each gene were left intact to preserve translatability of downstream genes when target genes were internal to a polycistronic message. Insertion sequences were matched to the transcriptional orientation of the original open reading frame. For *E. hormaechei,* the *acc(3)IV* apramycin resistance cassette was amplified from pUC18-miniTn7T-Apr (76) and used in place of kanamycin resistance. Recombination was facilitated by functions encoded on pKD46, pSIM19, or pSIM18 depending on the species mutated (74, 75). All mutations were confirmed by analyzing the sizes of PCR-amplified alleles and antibiotic resistance cassettes. In most cases mutations were further validated by sequencing. Recombineering plasmids were cured prior to phenotypic analysis. For selected mutants, complementation was achieved by either: a) cloning deleted gene sequences into pGEN-MCS (a plasmid stably maintained in these five type strains (77) during experimental infections in the absence of antibiotics) and transforming the resultant plasmid constructs into the respective mutants; or b) by allelic replacement into the chromosome with the wild-type gene.

### Bacterial growth rates

Bacterial strains were cultured in LB medium (78) with aeration and optical density (600 nm) was measured in 10-15 min intervals using either a LogPhase 600 (Agilent) or BioScreen C (Growth Curves USA) automated growth curve system. Maximal specific growth rates and doubling times were calculated using AMiGA software (17). Relative growth rates of each mutant were calculated for three biological replicates in comparison to a wild-type control cultured on the same microtiter plate.

### Murine model of bacteremia

Murine infections were performed in accordance with protocols approved by the University of Michigan Institutional Animal Care and Use Committee and were in accordance with Office of Laboratory Animal Welfare guidelines. Bacteria for murine infection experiments were prepared by subculturing overnight Luria broth growth into fresh medium and incubating for 2.5 hours. Exponential phase bacteria were collected by centrifugation and resuspended in an appropriate volume of PBS. Female 6–8-weeks old C57BL/6 mice were infected with bacterial suspension (containing the indicated number of cfu) of *E. coli* CFT073 (1 x 10^7^), *K. pneumoniae* KPPR1 (1 x 10^5^), S*. marcescens* UMH9 (5 x 10^6^), *C. freundii* UMH14 (1 x 10^8^), and *E. hormaechei* UM_CRE_14 (1 x 10^8^) via tail vein injection (79), unless otherwise noted. Total inocula for the five species were based on previous trials that 1) allowed for no bottleneck in the original trials (*i.e.*, no stochastic loss of mutants during the tail vein challenge); 2) accounted for the size and complexity of each transposon pool; and 3) avoided lethality following injection (see footnote ^a^ to Tables 3 and 4 for number of cfu in each inoculum). The spleen and liver from mice sacrificed at 24 hours post-inoculation were homogenized in PBS and ten-fold serial dilutions were plated on LB agar to determine the bacterial burden. For S. marcescens, kidneys were also used to determine bacterial burden. For competition infections, the wild-type *strain* was mixed with antibiotic-resistant mutant constructs at a 1:1 ratio prior to inoculation. The viable count for each strain was determined for both the inoculum (input) and organ homogenates (output) by serial dilution and differential plating on LB and LB containing antibiotics. The competitive index (CI) was calculated as follows: (CFUmutant/CFUwild-type)^output^/(CFUmutant/CFUwild-type)^input^. All murine infections were conducted using protocols approved by the University of Michigan Institutional Animal Care and Use Committee and in accordance with the Office of Laboratory Animal Welfare guidelines.

### Susceptibility to ciprofloxacin

Susceptibility to ciprofloxacin was measured by disk diffusion, which was performed for wild-type strains and their respective *xerC* and *ruvA* mutants. Luria broth was inoculated and cultured overnight at 37°C. Bacterial suspensions were normalized to an OD600 of 0.1 in accordance with the McFarland Standard Protocol (80). Sterile swabs were used to create a bacterial lawn on Mueller-Hinton agar plates. Ciprofloxacin disks (0.5 µg) were placed in the center of each agar plate using sterile forceps. Following incubation for 18-24 hours at 37°C, the diameter of the zone of inhibition was measured according to Kirby-Bauer Disk Diffusion Susceptibility Test Protocol (81).

### Measurement of proton motive force

To measure relative membrane potential, log-phase bacteria were diluted into PBS containing DiOC2 in DMSO. Fluorescence (red:green) was measured at excitation/emission of 483nm/501nm. CCCP was used to depolarize control samples as described (82).

### Susceptibility to antimicrobial peptides

CFU/ml and relative survival as compared to the wild-type strain survival was determined by incubating 10^7^ CFU/mL log-phase bacterial cells with polymyxin B in PBS, pH 7.4 for 45 min at 37°C and then plating for viable counts on Luria agar (82, 83). See legend to **Fig 3** for polymyxin B concentrations for each species.

### Susceptibility to human serum

Susceptibility to bactericidal activity of human serum was measured by incubating 10^7^ CFU/mL log-phase bacteria in 90% pooled human serum for *K. pneumoniae* KPPR1 and 40% pooled human serum for the other four species. Viability was measured by plating samples on Luria agar and determining CFUs after incubation times of 0 and 90 minutes. Both active serum and heat-inactivated (56°C, 60 min) serum were tested (10, 13, 84, 85).

### Siderophore production

Siderophore production was detected on chrome azurol S (CAS) plates supplemented with tryptone. Samples of overnight stationary phase cultures (2 µl) were spotted onto CAS agar plates and incubated at 37°C for 16 h. Siderophore activity was indicated by a shift from blue to yellow color within and surrounding the colonies. Gray scale image, which facilitate the detection of the blue (dark pixelation) to yellow (light pixelation) coloration in the CAS agar, was captured on a BioRad Gel Doc system with a “Coomassie Blue” setting and a white acrylic filter. The color image of the same plate was captured using an iPhone SE iOS 17.1.2. Media were prepared as described previously (86) with the sole modification of replacing 0.1% casamino acids with 1% tryptone to supplement the aromatic amino acid auxotroph inherent to mutation of *aroC*.

### Osmotic Stress

Susceptibility to osmotic stress was determined by incubating 10^7^ CFU/ml log-phase bacteria in PBS, pH7.4 with or without 2M D-sorbitol for 30 min at 37°C and then plating for viable counts (85, 87).

### Oxidative stress assessed by exposure to H_2_O_2_

To determine gene contributions to oxidative stress resistance, overnight bacterial cultures were adjusted to 1×10^7^ CFU/mL in PBS containing 1mM hydrogen peroxide. Each strain was incubated for 2 hours at 37°C, and quantitative culture was used to define the abundance of each strain at the input (t = 0) and output (t = 2) incubation. Percent survival was defined as [(CFU at t=2)/(CFU at t=0)] x 100. Fold change was defined as (wild-type %Survival/mutant %Survival) (8).

### Envelope stress

Envelope stress sensitivity was measured by growth on MacConkey agar, a medium containing bile salts (78). Mutants in *ruvA, tatC, gmhB*, and *wzxE* mutants and their respective wildtype strains in *E. coli, K. pneumoniae, C. freundii, S. marcescens*, were evaluated. Cultures were incubated at 37℃ overnight, then diluted in PBS to a final concentration of 10^4^ CFU/mL. Diluted cultures were then spread plated on MacConkey agar and LB agar in triplicate and incubated overnight at 27℃. Colonies were counted after 48 hours of incubation. The CFU on MacConkey agar was divided by the CFU on LB agar on each day and compared to the ratio of their respective WT for each day.

### Phosphate import

Phosphate transport mutants constitutively express alkaline phosphatase, whereas wild-type strains do not. Bacterial strains were cultured in phosphate-limiting minimal medium. Bacterial suspensions were normalized to OD600=0.1. Enzyme activity was measured by following hydrolysis of 0.4% *p*-nitrophenylphosphate at OD405 following incubation at 37°C for 1 hr as described (36, 78, 88).

### Twin Arginine protein export

To generate the *S. marcescens* SufI signal peptide-GFP translational fusion, the sequence encoding amino acid residues 1 to 35 of the N-terminal end of BVG96_RS17270 (89) was PCR amplified using Q5 polymerase (NEB) and cloned via NEBuilder HiFi DNA Assembly (NEB) into plasmid pIDMv5K-J231000-Dasher-GFP-B1006 (unpublished, S. Cocioba), previously modified with the replacement of the kanamycin resistance gene with a gene encoding gentamycin resistance. The resultant plasmid was confirmed by sequencing and transformed into *S. marcescens* UMH9 and Δ*tatC*::*nptII* via electroporation. Bacteria harboring the SufI-GFP fusion or vector control plasmid were cultured to mid exponential growth phase then fixed with 4% paraformaldehyde for 20 min at 25°C, washed with an equal volume of PBS, and normalized to 1×10^9^ CFU. Bacteria were mounted with Vectashield anti-fade medium (Vector Laboratories, Inc), and observed with a Nikon Ti2 Widefield microscope with a 100X oil immersion lens. Images were captured with an ORCA-Fusion Digital CMPS camera (Hamamatsu) and analyzed with ImageJ (Version 1.54f) to quantify cell length and fluorescence intensity (n>100).

## Ethics statement

Murine infections were performed in accordance with protocols approved by the University of Michigan Institutional Animal Care and Use Committee and were in accordance with Office of Laboratory Animal Welfare guidelines.

## Supporting Information

**Supplemental Fig. 1.**
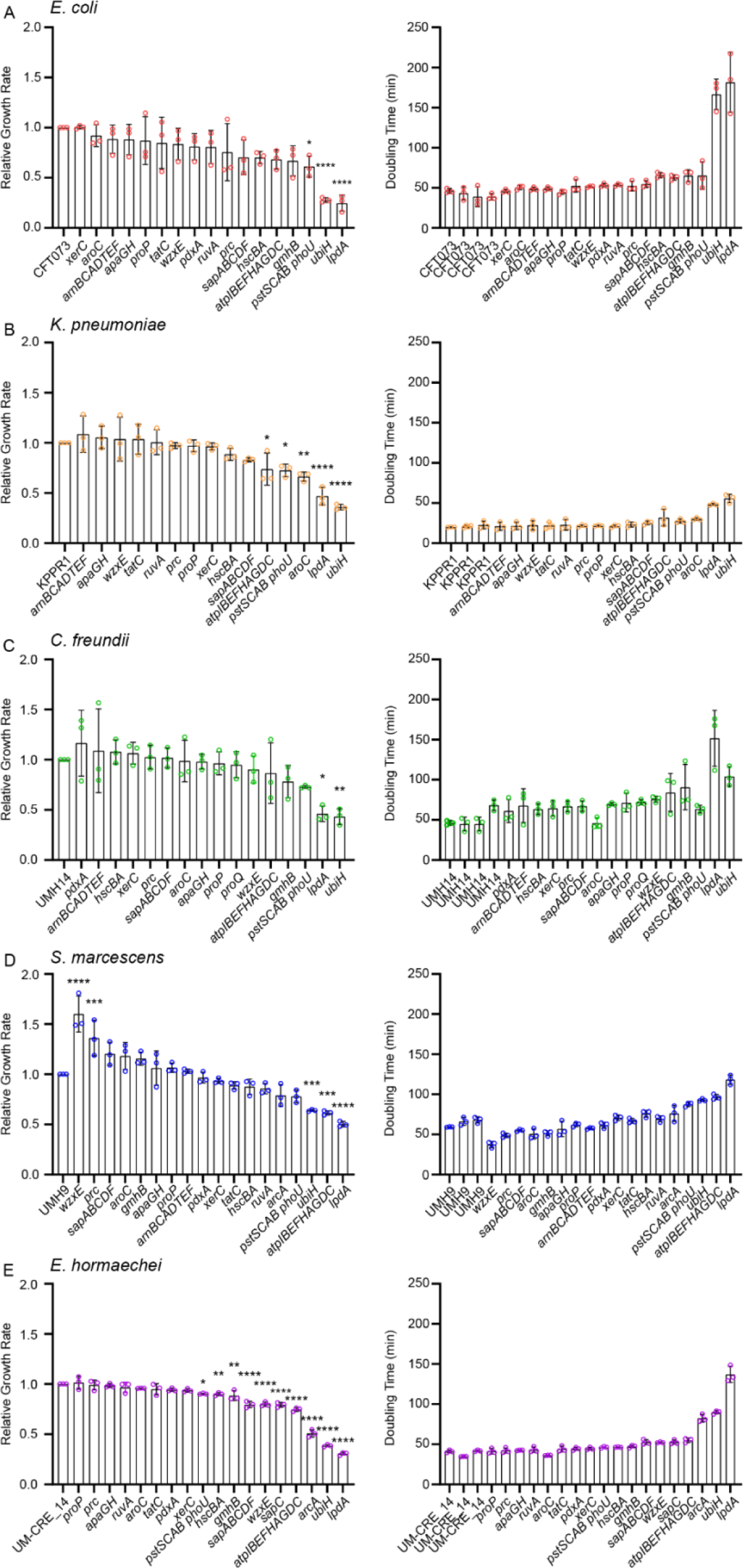
Growth characteristics of bacteremia fitness mutants. A-E. Relative growth rates (mutant/wild-type) and doubling times were calculated from the maximal specific growth rate in exponential phase for each mutant and wild-type strain. The statistical significance of relative growth rates was assessed by one-way ANOVA with Dunnett’s multiple comparisons test against the hypothetical value of 1 representing wild-type. Adjusted P values are indicated by asterisks: *, *p*<0.05; **, *p*<0.01; ***, *p*<0.001; ****, *p*<0.0001. Bars represent the means from three biological replicates ± the standard deviation. Wild-type controls from independent experimental plates were included for each species and set of mutants, as shown in doubling time calculations.

**Supplemental Fig 2.**
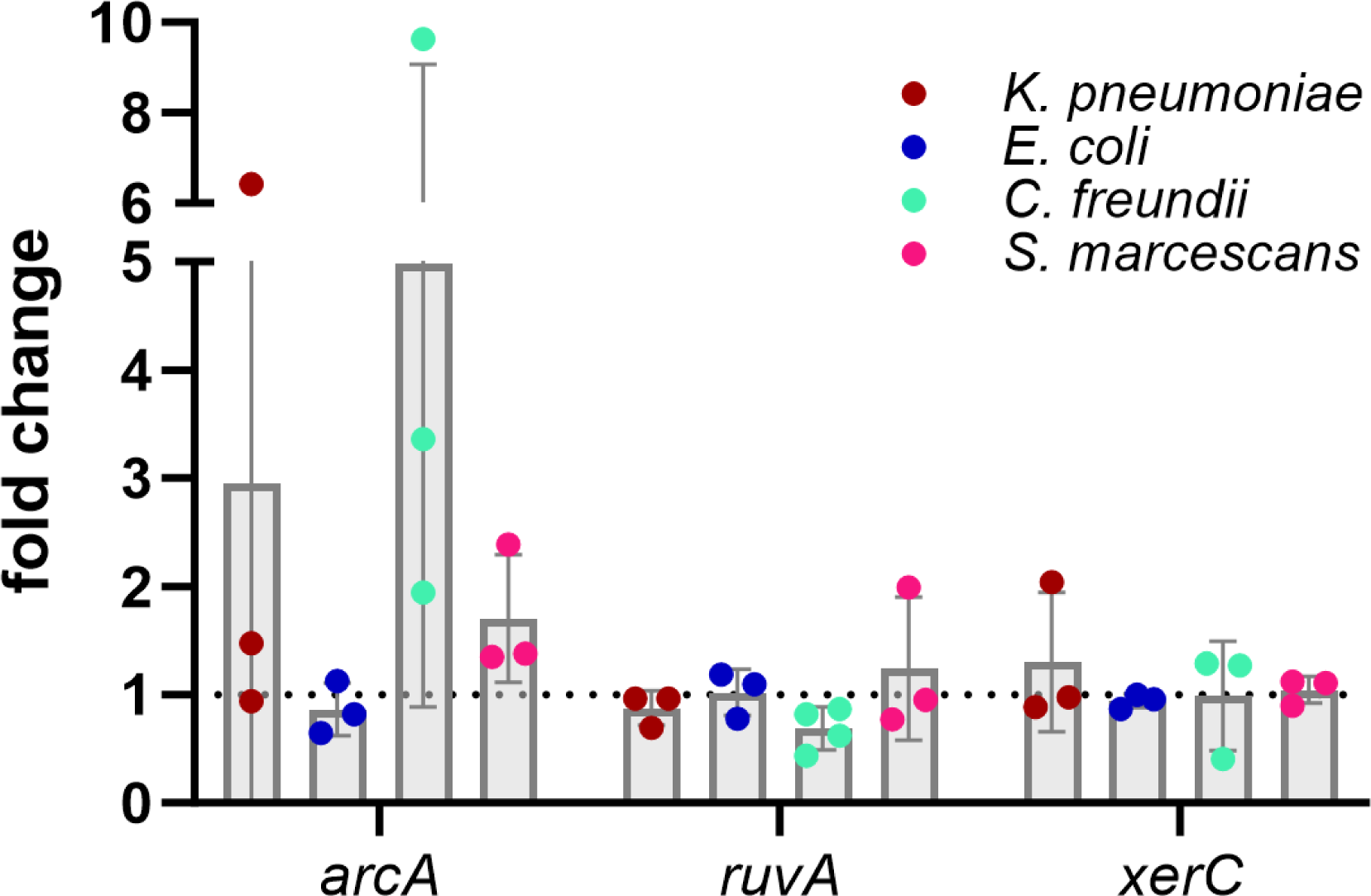
Oxidative stress by exposure to H_2_O_2._ Bacterial strains with mutations in the genes *arcA, ruvA,* and *xerC* in each species of interest were exposed to hydrogen peroxide to measure resistance to oxidative stress. Percent survival was calculated by comparing bacterial survival at 2 hours to the input. Fold change was calculated by dividing percent survival of each mutant to its respective wild-type strain. No mutant conveyed statistically significant resistance to hydrogen peroxide stress as assessed by a one-sample *t-*test with a hypothetical value of 1, representing survival of the wild-type strain. Data are means of 3 independent experiments.

**Supplemental Fig 3.**
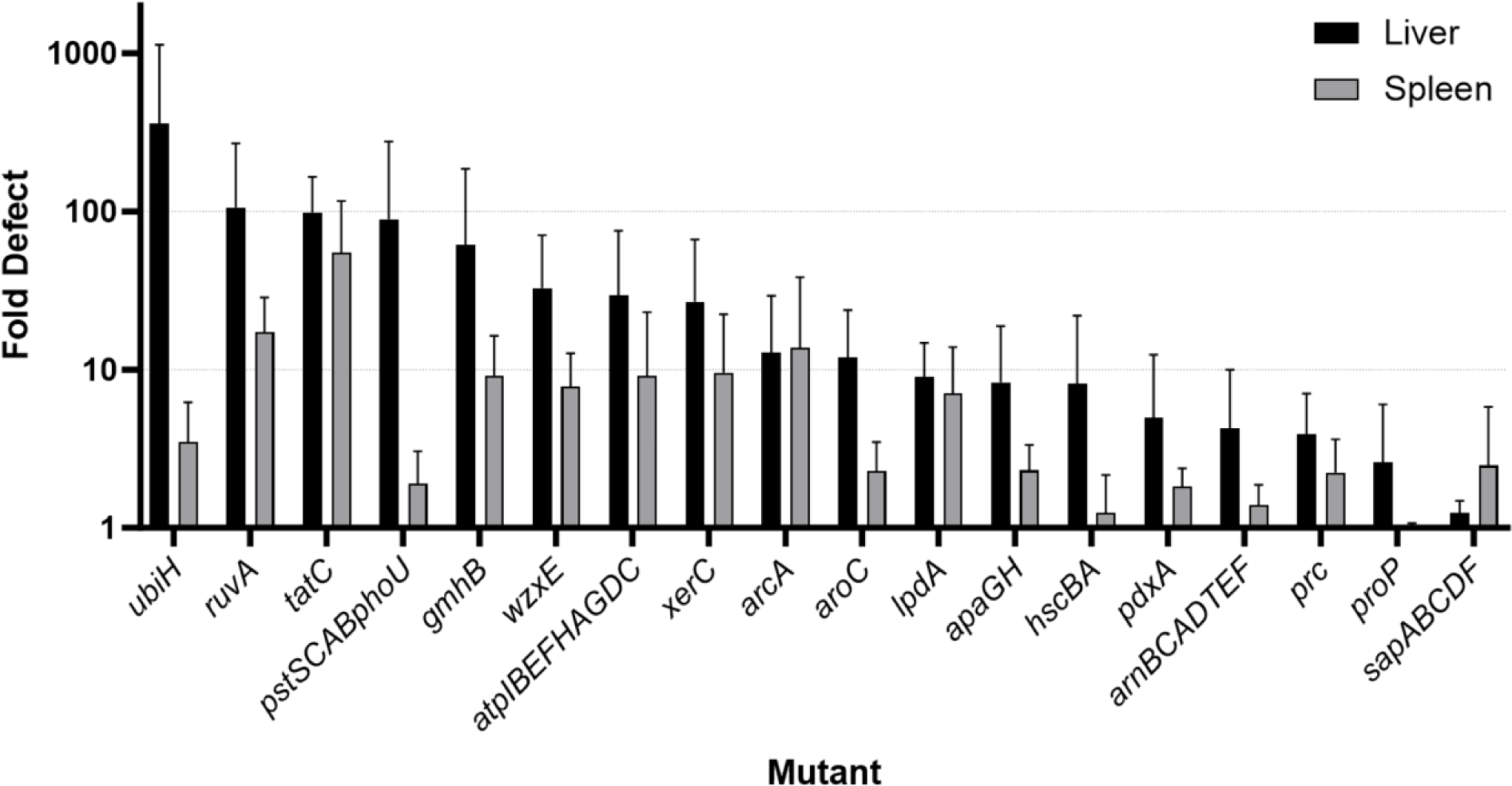
Fold-defects of fitness gene mutants as compared to wild-type strains in the murine model of bacteremia. Competitive indices in **Table 3** and **Table 4** have been converted to fold-defects, averaged for all species, and depicted separately for liver and spleen.

**Supplemental Table 1.**
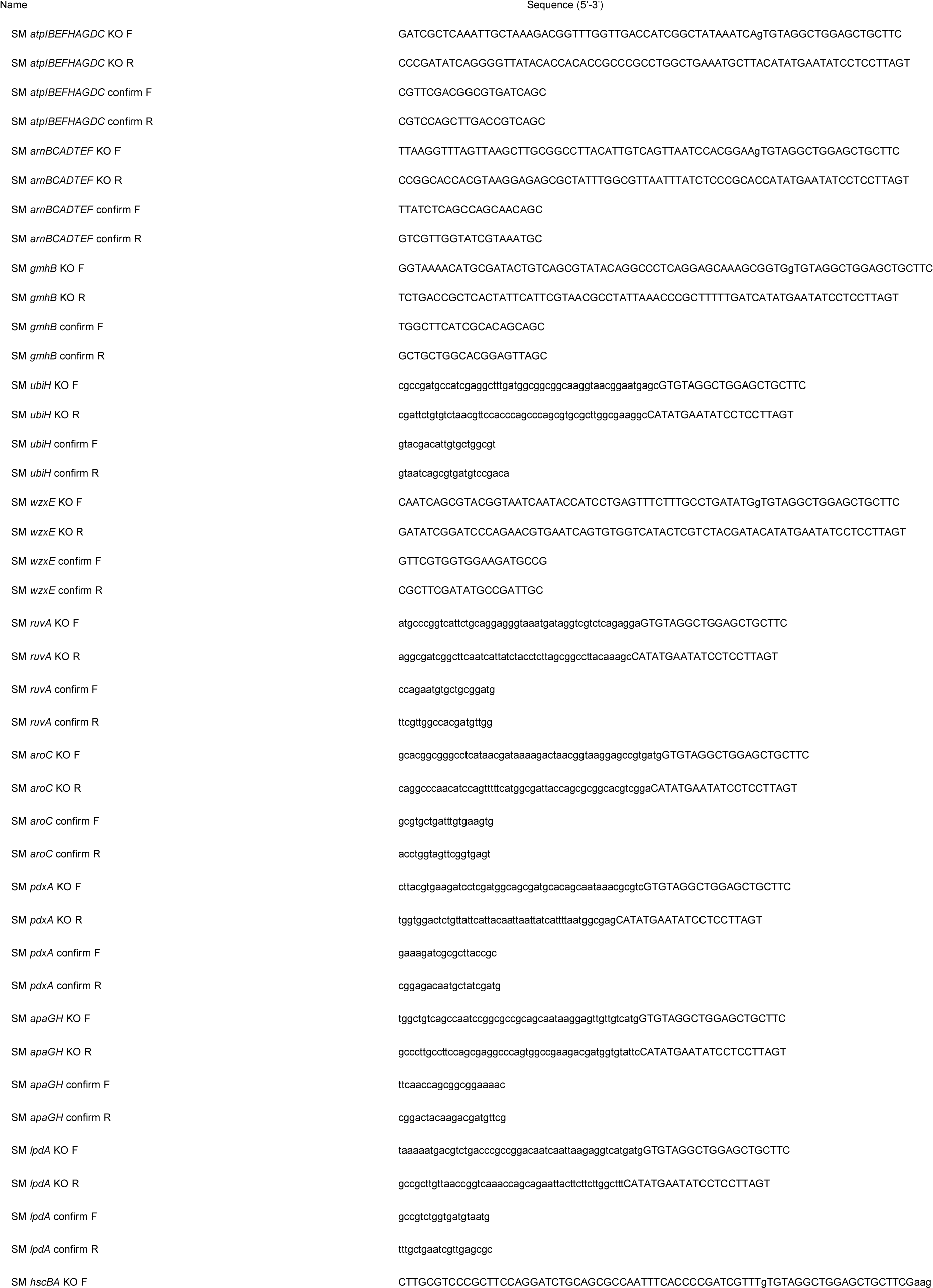
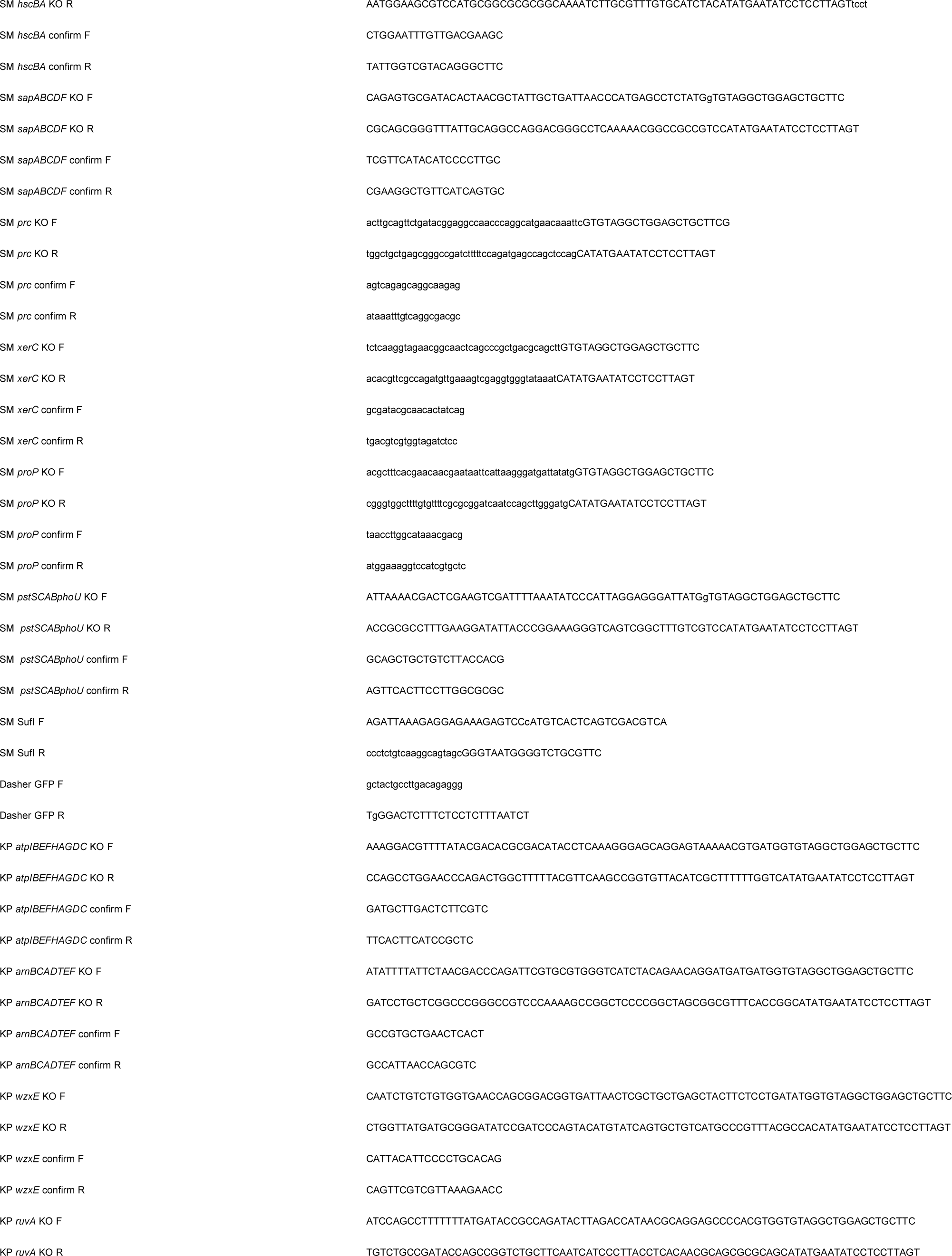
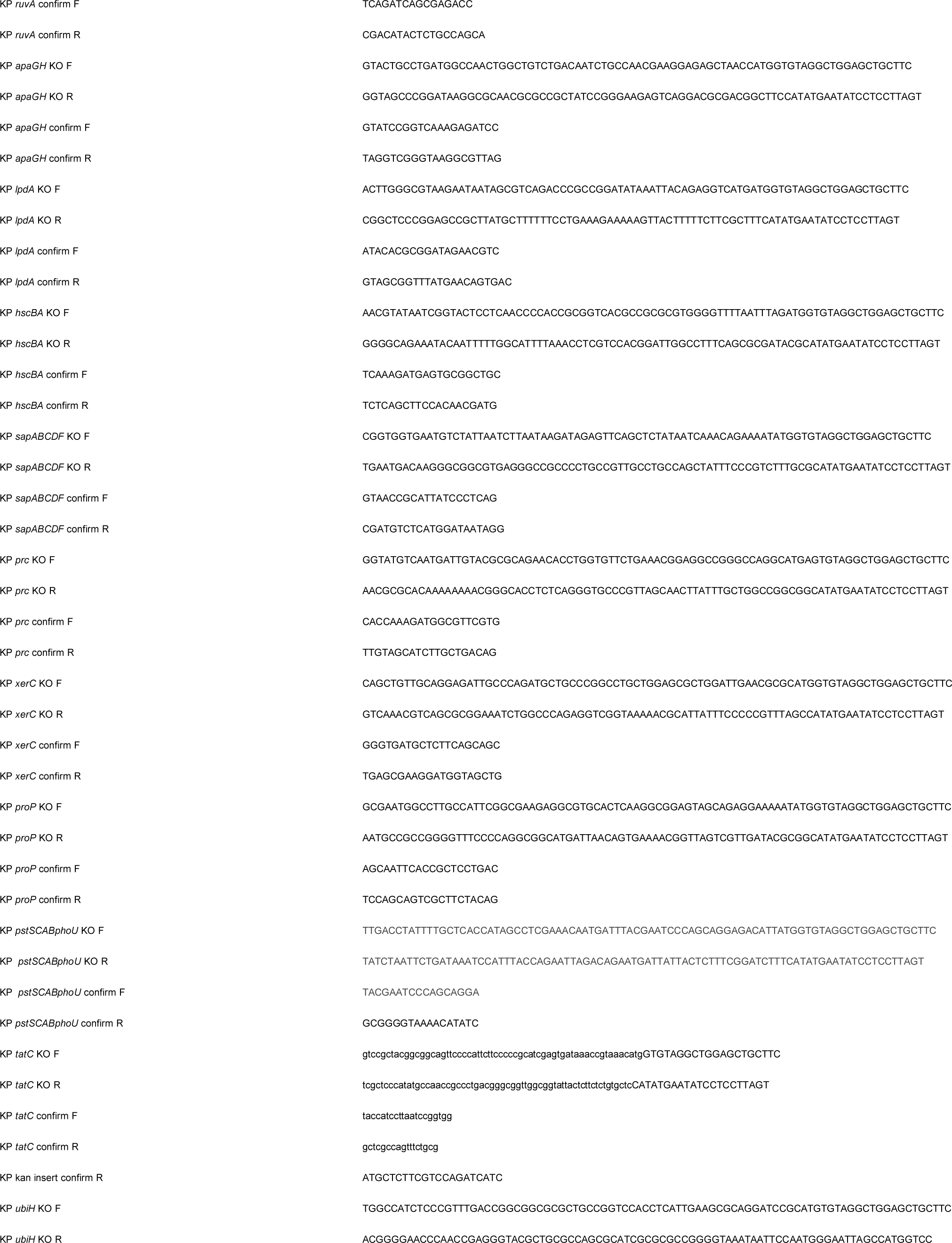
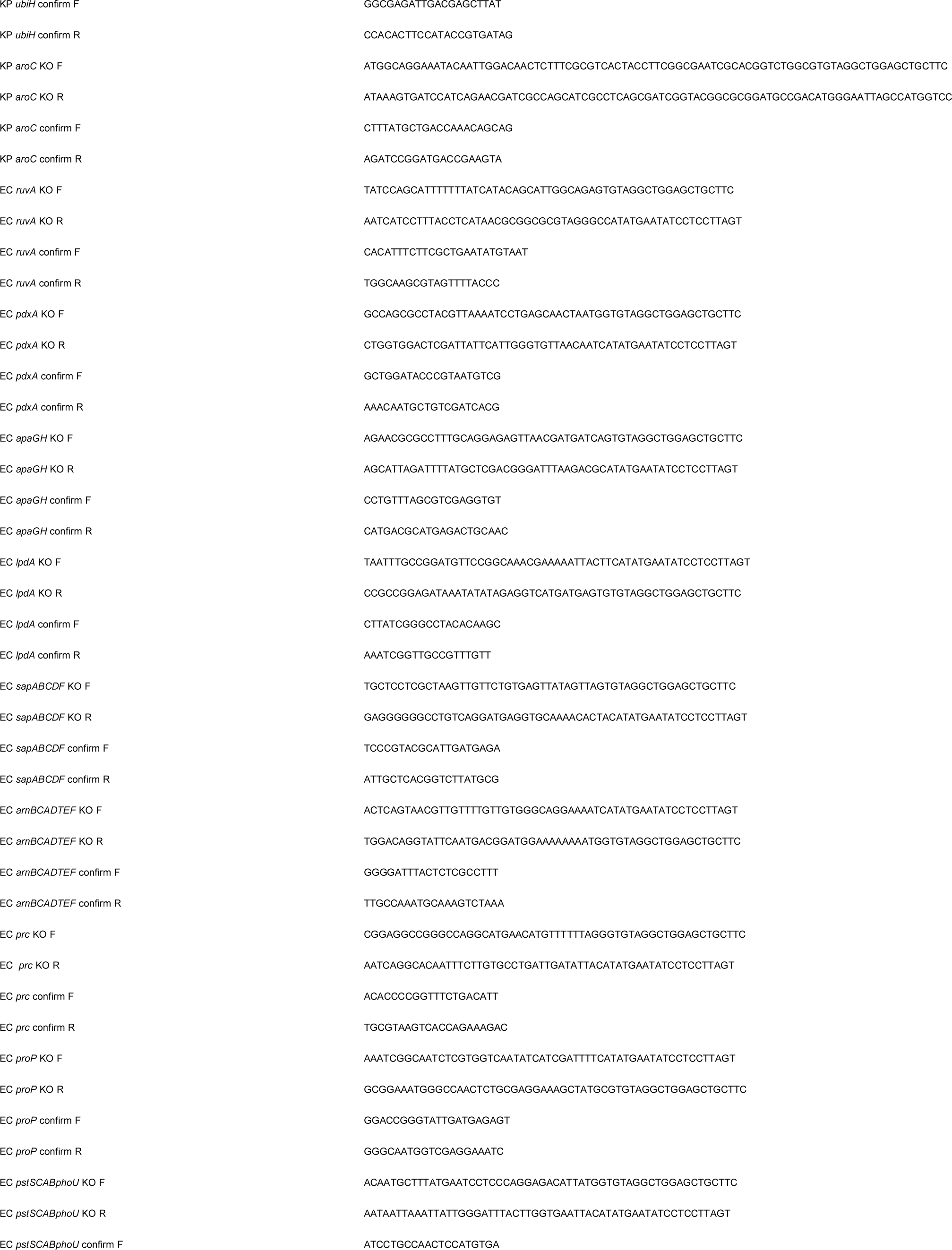
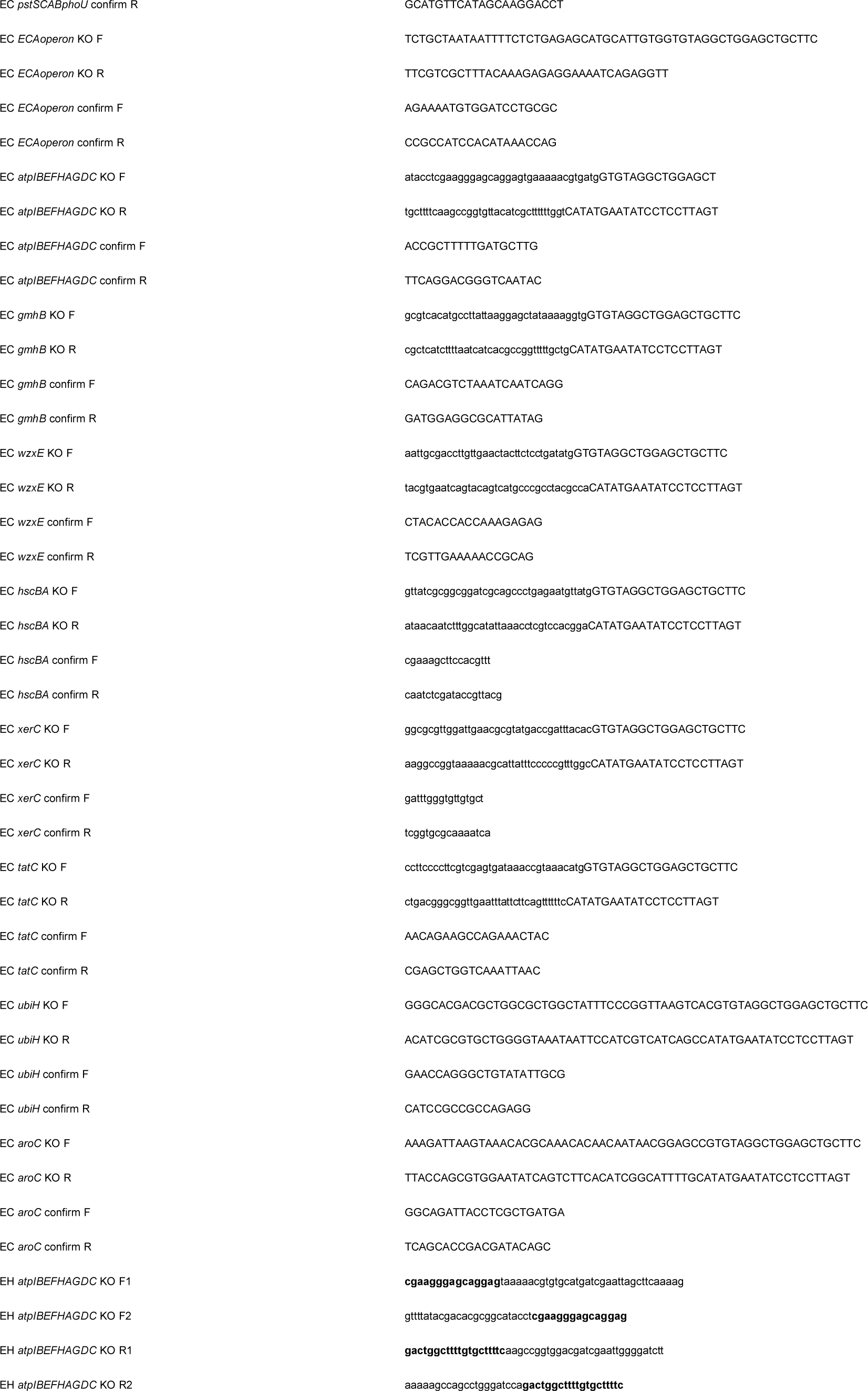
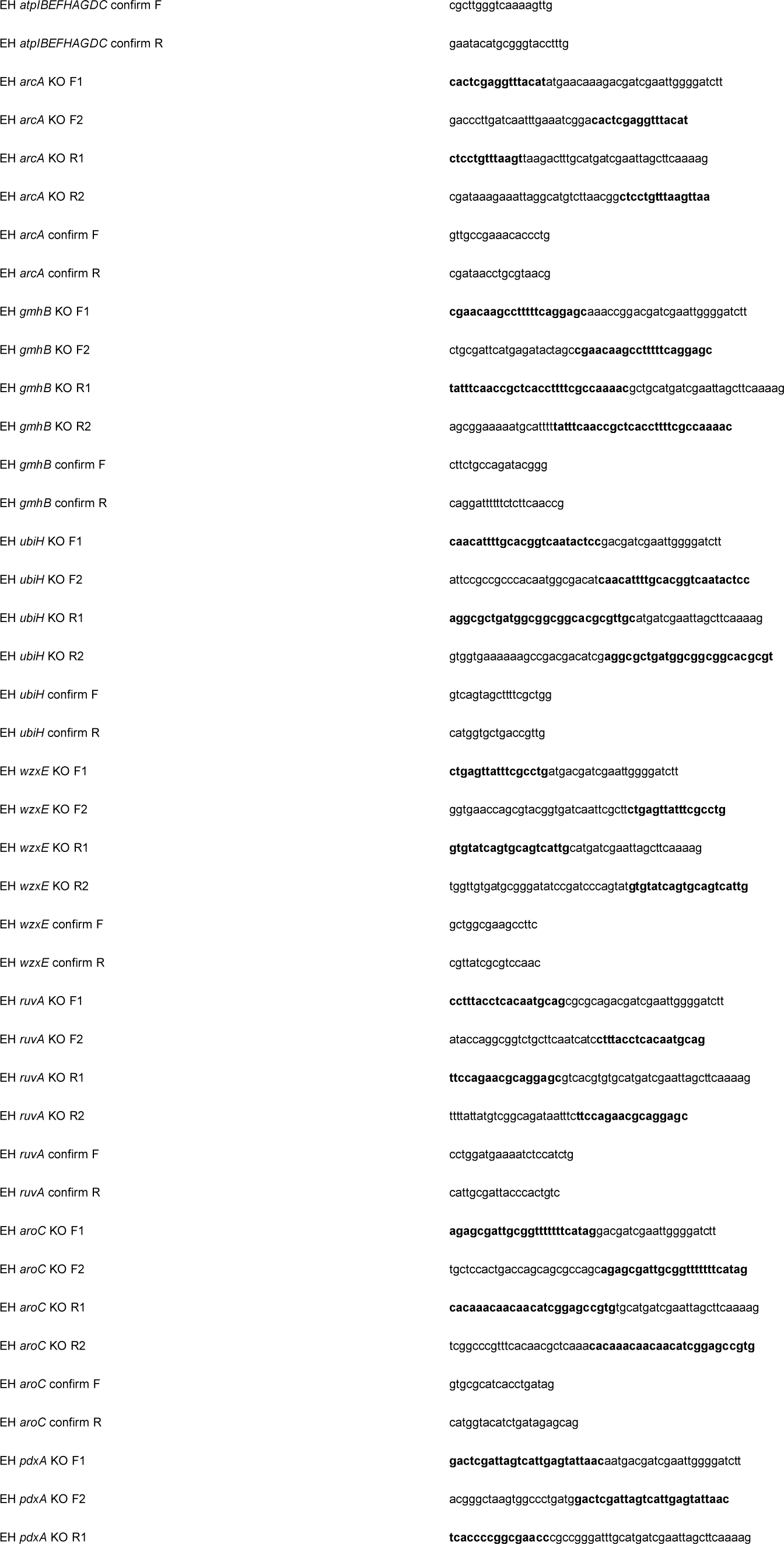
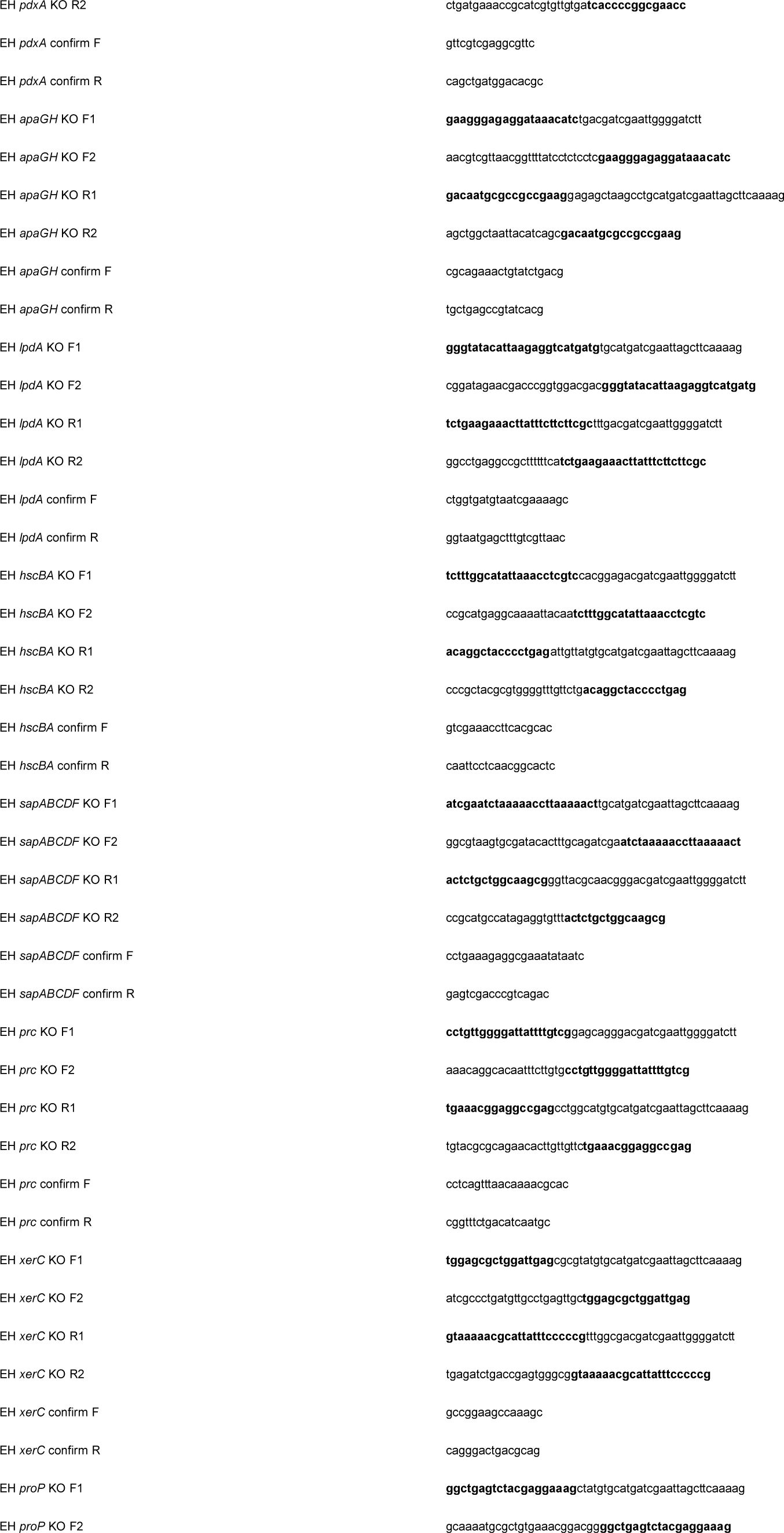
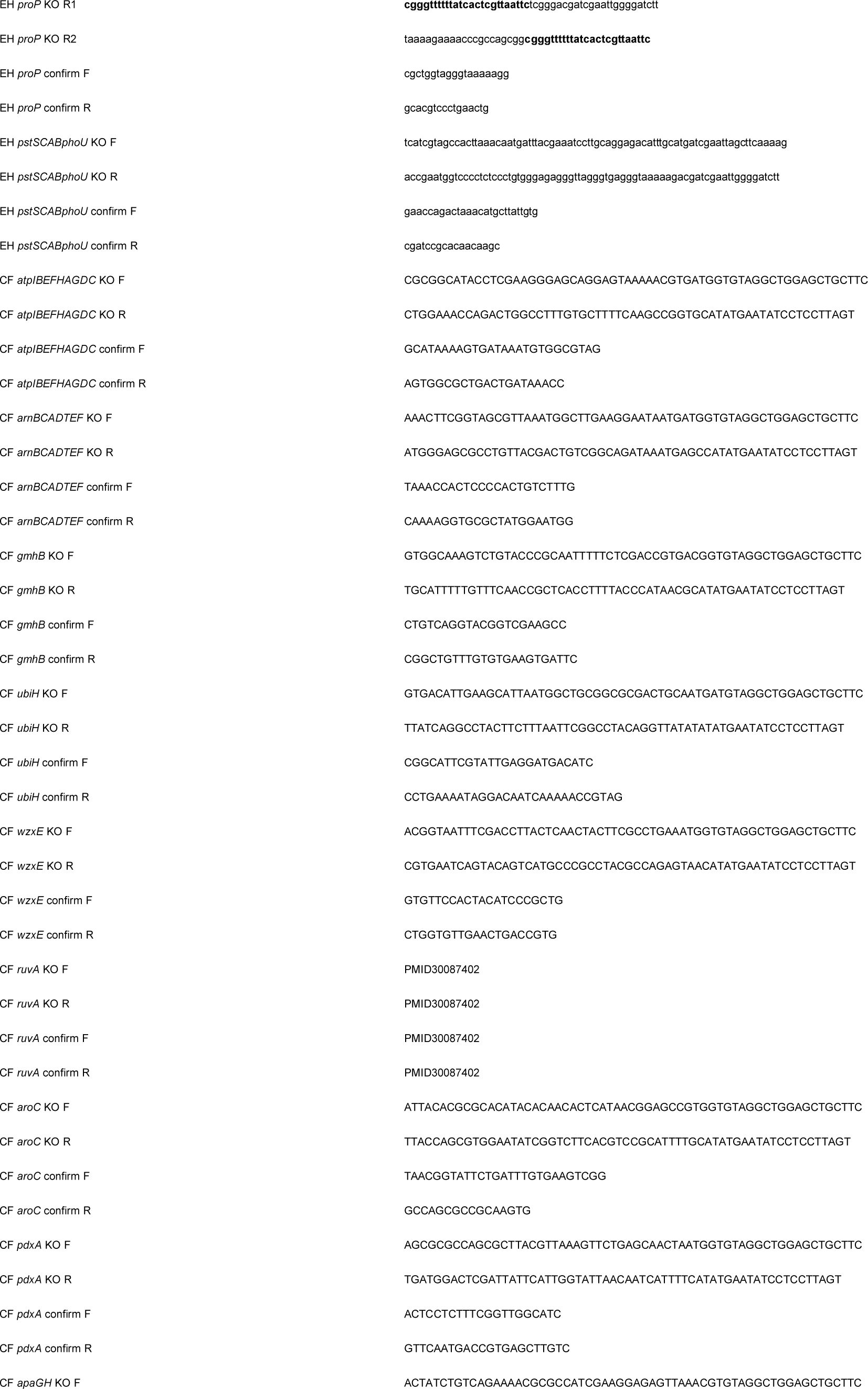
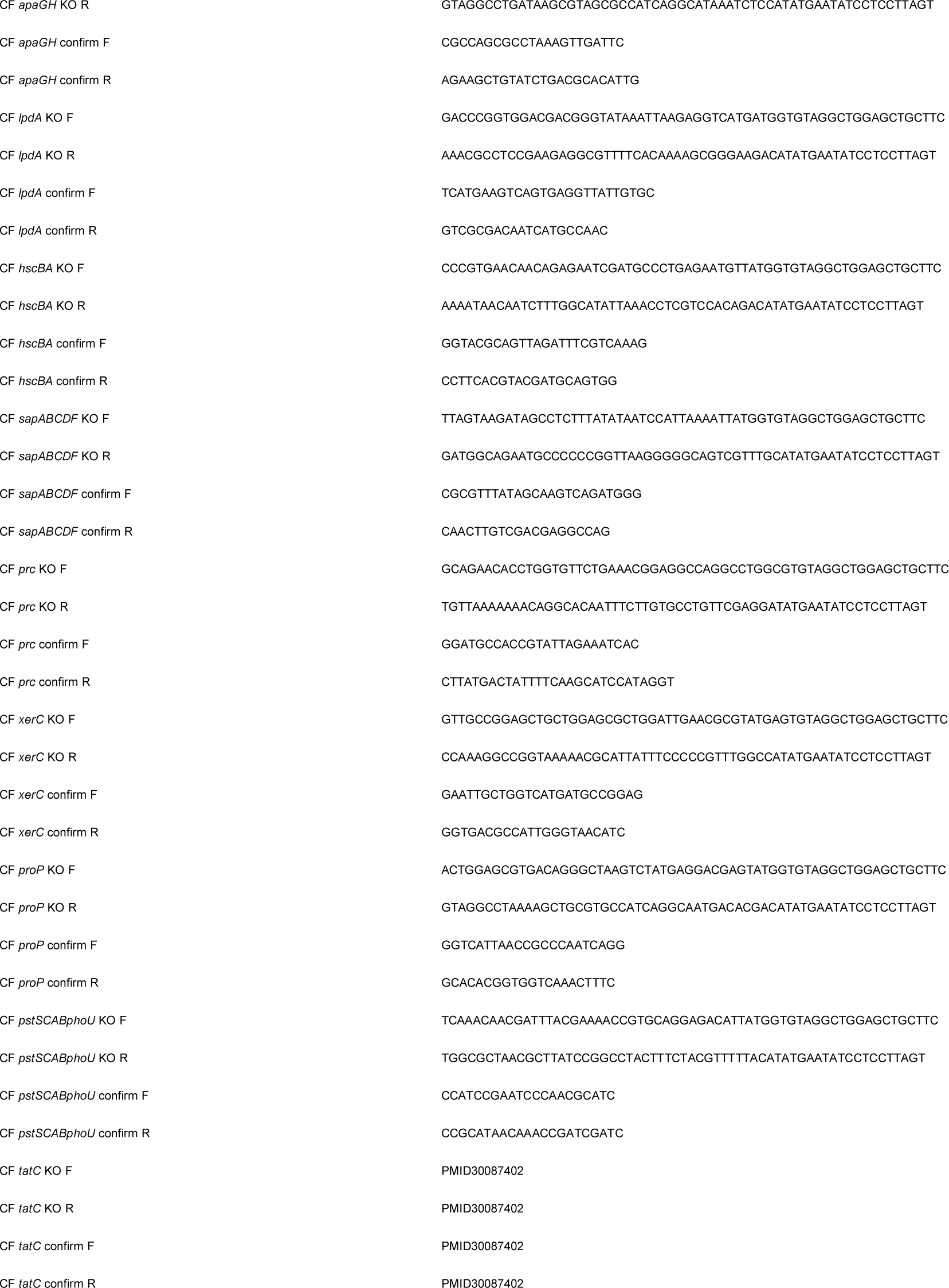
Primer sequences used for construction of prioritized fitness gene mutants.

**Supplemental Table 2.** Statistically significant *In vitro* phenotypes of fitness gene mutants.

| Mutant | *E. coli*<br>CFT073 | *K. pneumoniae*<br>KPPR1 | *S. marcescens*<br>UMH9 | *C. freundii*<br>UMH14 | *E. hormaechei*<br>UM_CRE_14 |
| --- | --- | --- | --- | --- | --- |
| *apaGH* |  |  |  |  |  |
| *arcA* | SC<sup>a</sup> | SC | SC | SC | SC |
| *arnBCADTEF* | PB | PB, LPS-M | PB, LPS-M | PB | <sub>-b</sub> |
| *aroC* | LowSid | LowSid | LowSid | LowSid |  |
| *atpIBEFHAGDC* |  |  |  |  |  |
| *gmhB* | BS |  |  |  |  |
| *hscBA* |  |  |  |  |  |
| *lpdA* |  |  |  |  |  |
| *pdxA* |  |  |  |  |  |
| *prc* | SS | SS | SS, OS | SS, OS | SS |
| *proP* |  |  |  |  |  |
| *pstSCABphoU* | AP | AP | AP | AP |  |
| *ruvA* | BS, CS | BS, CS | CS | CS | CS |
| *sapABCD* |  |  |  |  |  |
| *tatC* |  | BS | PL | BS |  |
| *ubiH* |  |  |  |  |  |
| *wzx* | SS, PB,<br>OS, BS | SS, OS | SS, PB, OS,<br>BS | PB, BS | SS, PB, OS |
| *xerC* | CS | CS | CS | CS |  |

<sup>a</sup>**Abbreviations:**
**SS**, human serum sensitivity; **PB**, polymyxin B sensitivity; **OS**, osmotic stress sensitivity; **BS**, bile salt sensitivity; **AP**, alkaline phosphatase overexpression; **LowSid**, mutation reduced siderophore activity; **SC**, small colony morphology; **CS**, ciprofloxacin susceptibility; **PL**, Tat-mediated protein localization; **LPS-M**, LPS modification.
<sup>b</sup>*arn* genes not present in *E. hormaechei* UMCRE14.

## References

1. Commission TJ. Sepsis 2022 [Available from: https://www.jointcommission.org/resources/patient-safety-topics/infection-prevention-and-control/sepsis/.

2. Diekema DJ, Hsueh PR, Mendes RE, Pfaller MA, Rolston KV, Sader HS, et al. The Microbiology of Bloodstream Infection: 20-Year Trends from the SENTRY Antimicrobial Surveillance Program. Antimicrob Agents Chemother. 2019;63(7).

3. Wisplinghoff H, Seifert H, Tallent SM, Bischoff T, Wenzel RP, Edmond MB. Nosocomial bloodstream infections in pediatric patients in United States hospitals: epidemiology, clinical features and susceptibilities. Pediatr Infect Dis J. 2003;22(8):686–91.

4. Wisplinghoff H, Bischoff T, Tallent SM, Seifert H, Wenzel RP, Edmond MB. Nosocomial bloodstream infections in US hospitals: analysis of 24,179 cases from a prospective nationwide surveillance study. Clin Infect Dis. 2004;39(3):309–17.

5. Holmes CL, Anderson MT, Mobley HLT, Bachman MA. Pathogenesis of Gram-Negative Bacteremia. Clin Microbiol Rev. 2021;34(2).

6. Anderson MT, Brown AN, Pirani A, Smith SN, Photenhauer AL, Sun Y, et al. Replication Dynamics for Six Gram-Negative Bacterial Species during Bloodstream Infection. mBio. 2021;12(4):e0111421.

7. Subashchandrabose S, Smith SN, Spurbeck RR, Kole MM, Mobley HL. Genome-wide detection of fitness genes in uropathogenic Escherichia coli during systemic infection. PLoS Pathog. 2013;9(12):e1003788.

8. Holmes CL, Wilcox AE, Forsyth V, Smith SN, Moricz BS, Unverdorben LV, et al. Klebsiella pneumoniae causes bacteremia using factors that mediate tissue-specific fitness and resistance to oxidative stress. PLoS Pathog. 2023;19(7):e1011233.

9. Mike LA, Stark AJ, Forsyth VS, Vornhagen J, Smith SN, Bachman MA, et al. A systematic analysis of hypermucoviscosity and capsule reveals distinct and overlapping genes that impact Klebsiella pneumoniae fitness. PLoS Pathog. 2021;17(3):e1009376.

10. Anderson MT, Mitchell LA, Zhao L, Mobley HLT. Capsule Production and Glucose Metabolism Dictate Fitness during Serratia marcescens Bacteremia. mBio. 2017;8(3).

11. Anderson MT, Mitchell LA, Zhao L, Mobley HLT. Citrobacter freundii fitness during bloodstream infection. Sci Rep. 2018;8(1):11792.

12. Subashchandrabose S, Hazen TH, Brumbaugh AR, Himpsl SD, Smith SN, Ernst RD, et al. Host-specific induction of Escherichia coli fitness genes during human urinary tract infection. Proc Natl Acad Sci U S A. 2014;111(51):18327–32.

13. Holmes CL, Smith SN, Gurczynski SJ, Severin GB, Unverdorben LV, Vornhagen J, et al. The ADP-Heptose Biosynthesis Enzyme GmhB is a Conserved Gram-Negative Bacteremia Fitness Factor. Infect Immun. 2022;90(7):e0022422.

14. Tamae C, Liu A, Kim K, Sitz D, Hong J, Becket E, et al. Determination of antibiotic hypersensitivity among 4,000 single-gene-knockout mutants of Escherichia coli. J Bacteriol. 2008;190(17):5981–8.

15. Liu A, Tran L, Becket E, Lee K, Chinn L, Park E, et al. Antibiotic sensitivity profiles determined with an Escherichia coli gene knockout collection: generating an antibiotic bar code. Antimicrob Agents Chemother. 2010;54(4):1393–403.

16. Brown AN, Anderson MT, Smith SN, Bachman MA, Mobley HLT. Conserved metabolic regulator ArcA responds to oxygen availability, iron limitation, and cell envelope perturbations during bacteremia. mBio. 2023;14(5):e0144823.

17. Midani FS, Collins J, Britton RA. AMiGA: Software for Automated Analysis of Microbial Growth Assays. mSystems. 2021;6(4):e0050821.

18. Breidenstein EB, Khaira BK, Wiegand I, Overhage J, Hancock RE. Complex ciprofloxacin resistome revealed by screening a Pseudomonas aeruginosa mutant library for altered susceptibility. Antimicrob Agents Chemother. 2008;52(12):4486–91.

19. Som N, Reddy M. Cross-talk between phospholipid synthesis and peptidoglycan expansion by a cell wall hydrolase. Proc Natl Acad Sci U S A. 2023;120(24):e2300784120.

20. Wang CY, Wang SW, Huang WC, Kim KS, Chang NS, Wang YH, et al. Prc contributes to Escherichia coli evasion of classical complement-mediated serum killing. Infect Immun. 2012;80(10):3399–409.

21. Parra-Lopez C, Baer MT, Groisman EA. Molecular genetic analysis of a locus required for resistance to antimicrobial peptides in Salmonella typhimurium. EMBO J. 1993;12(11):4053–62.

22. Stanley NR, Palmer T, Berks BC. The twin arginine consensus motif of Tat signal peptides is involved in Sec-independent protein targeting in Escherichia coli. J Biol Chem. 2000;275(16):11591–6.

23. Tarry M, Arends SJ, Roversi P, Piette E, Sargent F, Berks BC, et al. The Escherichia coli cell division protein and model Tat substrate SufI (FtsP) localizes to the septal ring and has a multicopper oxidase-like structure. J Mol Biol. 2009;386(2):504–19.

24. Stella NA, Callaghan JD, Zhang L, Brothers KM, Kowalski RP, Huang JJ, et al. SlpE is a calcium-dependent cytotoxic metalloprotease associated with clinical isolates of Serratia marcescens. Res Microbiol. 2017;168(6):567–74.

25. Berks BC. The twin-arginine protein translocation pathway. Annu Rev Biochem. 2015;84:843–64.

26. Bernhardt TG, de Boer PA. The Escherichia coli amidase AmiC is a periplasmic septal ring component exported via the twin-arginine transport pathway. Mol Microbiol. 2003;48(5):1171–82.

27. Ize B, Stanley NR, Buchanan G, Palmer T. Role of the Escherichia coli Tat pathway in outer membrane integrity. Mol Microbiol. 2003;48(5):1183–93.

28. Samaluru H, SaiSree L, Reddy M. Role of SufI (FtsP) in cell division of Escherichia coli: evidence for its involvement in stabilizing the assembly of the divisome. J Bacteriol. 2007;189(22):8044–52.

29. Crosa JH, Walsh CT. Genetics and assembly line enzymology of siderophore biosynthesis in bacteria. Microbiol Mol Biol Rev. 2002;66(2):223–49.

30. Weakland DR, Smith SN, Bell B, Tripathi A, Mobley HLT. The Serratia marcescens Siderophore Serratiochelin Is Necessary for Full Virulence during Bloodstream Infection. Infect Immun. 2020;88(8).

31. Bachman MA, Lenio S, Schmidt L, Oyler JE, Weiser JN. Interaction of lipocalin 2, transferrin, and siderophores determines the replicative niche of Klebsiella pneumoniae during pneumonia. mBio. 2012;3(6).

32. Bachman MA, Oyler JE, Burns SH, Caza M, Lepine F, Dozois CM, et al. Klebsiella pneumoniae yersiniabactin promotes respiratory tract infection through evasion of lipocalin 2. Infect Immun. 2011;79(8):3309–16.

33. Garcia EC, Brumbaugh AR, Mobley HL. Redundancy and specificity of Escherichia coli iron acquisition systems during urinary tract infection. Infect Immun. 2011;79(3):1225–35.

34. Wanner BL. Phosphorus assimilation and control of the phosphate regulon. In: Neidhardt FC, Curtiss, R. III, Ingraham, J. L., Lin, E. C. C., Low, K. B., Magasanik, B., Reznikoff, W. S., Riley, M., Schaechter, M., Umbarger, H. E., editor. Escherichia coli and Salmonella. Washington, DC: ASM Press; 1996. p. 1357-81.

35. Millan JL. Alkaline Phosphatases : Structure, substrate specificity and functional relatedness to other members of a large superfamily of enzymes. Purinergic Signal. 2006;2(2):335–41.

36. Jacobsen SM, Lane MC, Harro JM, Shirtliff ME, Mobley HL. The high-affinity phosphate transporter Pst is a virulence factor for Proteus mirabilis during complicated urinary tract infection. FEMS Immunol Med Microbiol. 2008;52(2):180–93.

37. Sha R, Liu F, Iwasaki H, Seeman NC. Parallel symmetric immobile DNA junctions as substrates for E. coli RuvC Holliday junction resolvase. Biochemistry. 2002;41(36):10985–93.

38. Huang WC, Lin CY, Hashimoto M, Wu JJ, Wang MC, Lin WH, et al. The role of the bacterial protease Prc in the uropathogenesis of extraintestinal pathogenic Escherichia coli. J Biomed Sci. 2020;27(1):14.

39. Rai AK, Mitchell AM. Enterobacterial Common Antigen: Synthesis and Function of an Enigmatic Molecule. mBio. 2020;11(4).

40. Arana DM, Ortega A, Gonzalez-Barbera E, Lara N, Bautista V, Gomez-Ruiz D, et al. Carbapenem-resistant Citrobacter spp. isolated in Spain from 2013 to 2015 produced a variety of carbapenemases including VIM-1, OXA-48, KPC-2, NDM-1 and VIM-2. J Antimicrob Chemother. 2017;72(12):3283–7.

41. Gonzalez LJ, Bahr G, Gonzalez MM, Bonomo RA, Vila AJ. In-cell kinetic stability is an essential trait in metallo-beta-lactamase evolution. Nat Chem Biol. 2023;19(9):1116–26.

42. Mason S, Vornhagen J, Smith SN, Mike LA, Mobley HLT, Bachman MA. The Klebsiella pneumoniae ter Operon Enhances Stress Tolerance. Infect Immun. 2023;91(2):e0055922.

43. Garcia-Weber D, Arrieumerlou C. ADP-heptose: a bacterial PAMP detected by the host sensor ALPK1. Cell Mol Life Sci. 2021;78(1):17–29.

44. Taylor PL, Sugiman-Marangos S, Zhang K, Valvano MA, Wright GD, Junop MS. Structural and kinetic characterization of the LPS biosynthetic enzyme D-alpha,beta-D-heptose-1,7-bisphosphate phosphatase (GmhB) from Escherichia coli. Biochemistry. 2010;49(5):1033–41.

45. Nikaido H. Molecular basis of bacterial outer membrane permeability revisited. Microbiol Mol Biol Rev. 2003;67(4):593–656.

46. Rollauer SE, Tarry MJ, Graham JE, Jaaskelainen M, Jager F, Johnson S, et al. Structure of the TatC core of the twin-arginine protein transport system. Nature. 2012;492(7428):210-4.

47. Palmer T, Berks BC. The twin-arginine translocation (Tat) protein export pathway. Nat Rev Microbiol. 2012;10(7):483–96.

48. O’May GA, Jacobsen SM, Longwell M, Stoodley P, Mobley HLT, Shirtliff ME. The high-affinity phosphate transporter Pst in Proteus mirabilis HI4320 and its importance in biofilm formation. Microbiology (Reading). 2009;155(Pt 5):1523–35.

49. Sinha R, LeVeque RM, Bowlin MQ, Gray MJ, DiRita VJ. Phosphate Transporter PstSCAB of Campylobacter jejuni Is a Critical Determinant of Lactate-Dependent Growth and Colonization in Chickens. J Bacteriol. 2020;202(7).

50. Dillingham MS, Kowalczykowski SC. RecBCD enzyme and the repair of double-stranded DNA breaks. Microbiol Mol Biol Rev. 2008;72(4):642–71, Table of Contents.

51. Wyatt HD, West SC. Holliday junction resolvases. Cold Spring Harb Perspect Biol. 2014;6(9):a023192.

52. Rice DW, Rafferty JB, Artymiuk PJ, Lloyd RG. Insights into the mechanisms of homologous recombination from the structure of RuvA. Curr Opin Struct Biol. 1997;7(6):798–803.

53. West SC. The RuvABC proteins and Holliday junction processing in Escherichia coli. J Bacteriol. 1996;178(5):1237–41.

54. Otsuji N, Iyehara H, Hideshima Y. Isolation and characterization of an Escherichia coli ruv mutant which forms nonseptate filaments after low doses of ultraviolet light irradiation. J Bacteriol. 1974;117(2):337–44.

55. Blakely G, Colloms S, May G, Burke M, Sherratt D. Escherichia coli XerC recombinase is required for chromosomal segregation at cell division. New Biol. 1991;3(8):789–98.

56. Ledger EVK, Lau K, Tate EW, Edwards AM. XerC Is Required for the Repair of Antibiotic- and Immune-Mediated DNA Damage in Staphylococcus aureus. Antimicrob Agents Chemother. 2023;67(3):e0120622.

57. Herrmann KM, Weaver LM. The Shikimate Pathway. Annu Rev Plant Physiol Plant Mol Biol. 1999;50:473–503.

58. Cox GB, Gibson F. Biosynthesis of Vitamin K and Ubiquinone. Relation to the Shikimic Acid Pathway in Escherichia Coli. Biochim Biophys Acta. 1964;93:204–6.

59. Bachman MA, Breen P, Deornellas V, Mu Q, Zhao L, Wu W, et al. Genome-Wide Identification of Klebsiella pneumoniae Fitness Genes during Lung Infection. mBio. 2015;6(3):e00775.

60. Karki HS, Ham JH. The roles of the shikimate pathway genes, aroA and aroB, in virulence, growth and UV tolerance of Burkholderia glumae strain 411gr-6. Mol Plant Pathol. 2014;15(9):940–7.

61. Hernanz Moral C, Flano del Castillo E, Lopez Fierro P, Villena Cortes A, Anguita Castillo J, Cascon Soriano A, et al. Molecular characterization of the Aeromonas hydrophila aroA gene and potential use of an auxotrophic aroA mutant as a live attenuated vaccine. Infect Immun. 1998;66(5):1813–21.

62. Felgner S, Frahm M, Kocijancic D, Rohde M, Eckweiler D, Bielecka A, et al. aroA-Deficient Salmonella enterica Serovar Typhimurium Is More Than a Metabolically Attenuated Mutant. mBio. 2016;7(5).

63. Wallace BJ, Young IG. Role of quinones in electron transport to oxygen and nitrate in Escherichia coli. Studies with a ubiA-menA-double quinone mutant. Biochim Biophys Acta. 1977;461(1):84–100.

64. Young IG, Stroobant P, Macdonald CG, Gibson F. Pathway for ubiquinone biosynthesis in Escherichia coli K-12: gene-enzyme relationships and intermediates. J Bacteriol. 1973;114(1):42–52.

65. Meganathan R, Kwon O. Biosynthesis of Menaquinone (Vitamin K2) and Ubiquinone (Coenzyme Q). EcoSal Plus. 2009;3(2).

66. Malpica R, Sandoval GR, Rodriguez C, Franco B, Georgellis D. Signaling by the arc two-component system provides a link between the redox state of the quinone pool and gene expression. Antioxid Redox Signal. 2006;8(5-6):781–95.

67. Brown AN, Anderson MT, Bachman MA, Mobley HLT. The ArcAB Two-Component System: Function in Metabolism, Redox Control, and Infection. Microbiol Mol Biol Rev. 2022;86(2):e0011021.

68. Thaden JT, Li Y, Ruffin F, Maskarinec SA, Hill-Rorie JM, Wanda LC, et al. Increased Costs Associated with Bloodstream Infections Caused by Multidrug-Resistant Gram-Negative Bacteria Are Due Primarily to Patients with Hospital-Acquired Infections. Antimicrob Agents Chemother. 2017;61(3):e01709–16.

69. Morris S, Cerceo E. Trends, Epidemiology, and Management of Multi-Drug Resistant Gram-Negative Bacterial Infections in the Hospitalized Setting. Antibiotics (Basel). 2020;9(4).

70. Kang CI, Kim SH, Park WB, Lee KD, Kim HB, Kim EC, et al. Bloodstream infections caused by antibiotic-resistant gram-negative bacilli: risk factors for mortality and impact of inappropriate initial antimicrobial therapy on outcome. Antimicrob Agents Chemother. 2005;49(2):760–6.

71. Andre AC, Debande L, Marteyn BS. The selective advantage of facultative anaerobes relies on their unique ability to cope with changing oxygen levels during infection. Cell Microbiol. 2021;23(8):e13338.

72. Subashchandrabose S, Smith S, DeOrnellas V, Crepin S, Kole M, Zahdeh C, et al. Acinetobacter baumannii Genes Required for Bacterial Survival during Bloodstream Infection. mSphere. 2016;1(1).

73. Pont S, Janet-Maitre M, Faudry E, Cretin F, Attree I. Molecular Mechanisms Involved in Pseudomonas aeruginosa Bacteremia. Adv Exp Med Biol. 2022;1386:325–45.

74. Datsenko KA, Wanner BL. One-step inactivation of chromosomal genes in Escherichia coli K-12 using PCR products. Proc Natl Acad Sci U S A. 2000;97(12):6640–5.

75. Datta S, Costantino N, Court DL. A set of recombineering plasmids for gram-negative bacteria. Gene. 2006;379:109–15.

76. Ducas-Mowchun K, De Silva PM, Crisostomo L, Fernando DM, Chao TC, Pelka P, et al. Next Generation of Tn7-Based Single-Copy Insertion Elements for Use in Multi- and Pan-Drug-Resistant Strains of Acinetobacter baumannii. Appl Environ Microbiol. 2019;85(11).

77. Lane MC, Alteri CJ, Smith SN, Mobley HL. Expression of flagella is coincident with uropathogenic Escherichia coli ascension to the upper urinary tract. Proc Natl Acad Sci U S A. 2007;104(42):16669–74.

78. Miller JH. Experiments in Molecular Genetics. Cold Spring Harbor, N. Y.: Cold Spring Harbor Laboratory; 1972.

79. Smith SN, Hagan EC, Lane MC, Mobley HL. Dissemination and systemic colonization of uropathogenic Escherichia coli in a murine model of bacteremia. mBio. 2010;1(5).

80. Clinical Microbiology Procedures Handbook. 4th ed. Washington DC: ASM Press; 2016.

81. Hudzicki J. Kirby-Bauer Disk Diffusion Susceptibility Test Protocol: ASM Press; 2009 [Available from: https://asm.org/getattachment/2594ce26-bd44-47f6-8287-0657aa9185ad/Kirby-Bauer-Disk-Diffusion-Susceptibility-Test-Protocol-pdf.pdf.

82. Alteri CJ, Lindner JR, Reiss DJ, Smith SN, Mobley HL. The broadly conserved regulator PhoP links pathogen virulence and membrane potential in Escherichia coli. Mol Microbiol. 2011;82(1):145–63.

83. Meier-Dieter U, Acker G, Mayer H. Detection of enterobacterial common antigen on bacterial cell surfaces by colony-immunoblotting: effect of its linkage to lipopolysaccharide. FEMS Microbiol Lett. 1989;50(1-2):215–9.

84. Anderson MT, Himpsl SD, Mitchell LA, Kingsley LG, Snider EP, Mobley HLT. Identification of distinct capsule types associated with Serratia marcescens infection isolates. PLoS Pathog. 2022;18(3):e1010423.

85. Shea AE, Frick-Cheng AE, Smith SN, Mobley HLT. Phenotypic Assessment of Clinical Escherichia coli Isolates Predicts Uropathogenic Potential. mSystems. 2022:e0082722.

86. Himpsl SD, Mobley HLT. Siderophore Detection Using Chrome Azurol S and Cross-Feeding Assays. Methods Mol Biol. 2019;2021:97–108.

87. Rojas E, Theriot JA, Huang KC. Response of Escherichia coli growth rate to osmotic shock. Proc Natl Acad Sci U S A. 2014;111(21):7807–12.

88. Wanner BL. Novel regulatory mutants of the phosphate regulon in Escherichia coli K-12. J Mol Biol. 1986;191(1):39–58.

89. Bendtsen JD, Nielsen H, Widdick D, Palmer T, Brunak S. Prediction of twin-arginine signal peptides. BMC Bioinformatics. 2005;6:167.

